# The function and decline of the female reproductive tract at single-cell resolution

**DOI:** 10.1101/2022.10.26.513823

**Authors:** Ivana Winkler, Alexander Tolkachov, Fritjof Lammers, Perrine Lacour, Nina Schneider, Marie-Luise Koch, Jasper Panten, Florian Grünschläger, Klaudija Daugelaite, Tanja Poth, Simon Haas, Duncan T. Odom, Angela Goncalves

## Abstract

The female reproductive tract (FRT) undergoes extensive remodeling during each reproductive cycle, regulated by systemic changes in sex hormones. Whether this recurrent remodeling influences a specific organ’s aging trajectory is unknown. To address this, we systematically characterized at single-cell resolution the morphological and transcriptional changes that occur in ovary, oviduct, uterus, cervix, and vagina at each phase of the mouse estrus cycle, during decidualization, and into aging. Transcriptional and cell-to-cell communication networks in estrus cycle and aging are enriched for ECM reorganization and inflammation, two essential components of FRT remodeling. We directly link the organ-specific level of these two processes over reproductive lifespan with the gradual, age-related development of fibrosis and chronic inflammation. Our data represent a comprehensive atlas of the FRT lifespan, revealing pathological consequences of incomplete resolution of recurrent inflammation and tissue repair.

## Introduction

During the estrus cycle the mammalian female reproductive tract (FRT) undergoes extensive remodeling in preparation for ovulation and pregnancy. The physiological changes to the FRT, which occur in response to ovarian steroid hormones, are conserved between humans and other mammals, with the exception of the human-specific spontaneous terminal differentiation of the endometrial stromal cells in the process of decidualization (Bellofiore et al., 2018). These decidualized stromal cells are subsequently expelled during human menstruation. In most other mammals this final step of differentiation can be modeled by pregnancy induction (Miller and Takahashi, 2014; Rajkovic et al., 2004). The reproductive cycle influences important functions of other organ systems, including shaping the immune response to infection and neural plasticity (Fernández et al., 2003; Gallichan and Rosenthal, 1996), yet remains poorly characterized. Prior analyses of mammalian estrus cycle have had a number of limitations: they were often microscopy-based (Garry et al., 2010; Hickey et al., 2013; Jurgensen et al., 1996; Sato et al., 1997; Schulke et al., 2008; Wang et al., 2000), analyzed single organs of the FRT (Garcia-Alonso et al., 2021; Jemt et al., 2016; Roberson et al., 2021; Saare et al., 2016; Wang et al., 2020), assayed the activity of few genes (Cornet et al., 2002; Von Wolff et al., 1999), used bulk tissues (Kim et al., 2018), and/or have been primarily qualitative (Igarashi et al., 1995).

Although the FRT organs are regulated systemically by sex hormones, they vary extensively in their susceptibility to age-related pathologies. Development of chronic degenerative diseases during aging is triggered by numerous changes in the cell microenvironment and cell-to-cell interactions. Systemic inflammation, known as inflammaging, significantly contributes to age-related morbidities. Inflammation is often accompanied by excessive accumulation of extracellular matrix (ECM), resulting in fibrosis. Progressive development of fibrosis with aging can lead to organ function impairment (Horowitz and Thannickal, 2019).

In mammals, many conserved reproductive processes, including ovulation in the ovary, menstruation and decidualization/implantation in the uterus, and the remodeling of the vaginal epithelium throughout the estrus cycle, display hallmark signs of inflammation (Jabbour et al., 2009) and ECM remodeling (Salamonsen et al., 2002). The female reproductive tract can resolve these cyclical inflammatory events rapidly and thus re-establish normal reproductive function. Multiple cell types cooperate to fine-tune the complex process of inflammation, with immune cells and fibroblasts playing major roles. Immune cells recognize and eliminate inflammatory triggers, while fibroblasts reorganize the microenvironment through expression of inflammatory cytokines, extracellular matrix (ECM) components and remodeling enzymes. When inflammation is not normally resolved, because of aging or other factors, chronic inflammation and fibrosis can develop. Fibroblasts likely shape inflammation persistence (Davidson et al., 2021) by failing to return to a homeostatic state, thereby contributing to inflammatory memory (Kirk et al., 2021). As the main ECM producers, they play key roles in tissue remodeling. Excessive and/or frequent tissue remodeling as a consequence of reoccurring injury can lead to fibrosis development (Rockey et al., 2015), but current models assume that the FRT cyclical remodeling is scar-free.

To systematically explore the female reproductive cycle, we characterized at single-cell resolution the morphological and transcriptional changes that occur in ovary, oviduct, uterus, cervix, and vagina at each phase of the mouse estrus cycle, during decidualization, and into old age. Specifically to enable inter-organ comparisons at the same cycle stages, all five FRT organs were simultaneously collected and analyzed from over 20 individual mice. Our analyses newly reveal how the physiological differences between the upper (ovary, oviduct, uterus) and lower reproductive tract (cervix, vagina) are closely mirrored by compositional and transcriptional differences. To explore whether the cyclic inflammation and remodeling that naturally occur during the reproductive lifespan of young mice result in age-related chronic inflammation and fibrosis, we extensively characterized the inflammatory status of fibroblasts and their cell-to-cell communication networks during normal cycling and aging. We determined that transcription factor and cell-to-cell communication networks active in fibroblasts during estrus cycling and aging are enriched for ECM remodeling and inflammation, and are conserved between humans and mouse uteruses. Our analysis reveals that re-occurring cyclic changes in inflammation and ECM activity in each reproductive cycle significantly contribute to FRT aging linking the number of elapsed cycles with the age-related accumulation of fibrosis and inflammation.

## Results

### Single-cell characterization of the cycling female reproductive tract

We profiled how the female reproductive tract is remodeled during the estrus cycle, decidualization, and aging using single-cell RNA-sequencing (scRNA-seq). We characterized the cellular composition and transcriptional states present in ovary, oviduct, uterus, cervix, and vagina (with spleen as a control organ) by collecting 378,516 single-cell transcriptomes from normally cycling young mice in each of the four cycle phases (P - proestrus, E - estrus, M - metestrus and D - diestrus), as well as 30,966 cells from early pregnancy and 74,129 cells from acyclic old mice (Figure 1a-b, Table S1). All experiments were performed in three to seven biological replicates to allow for the assessment of statistical significance in all comparisons.

**Figure 1.**
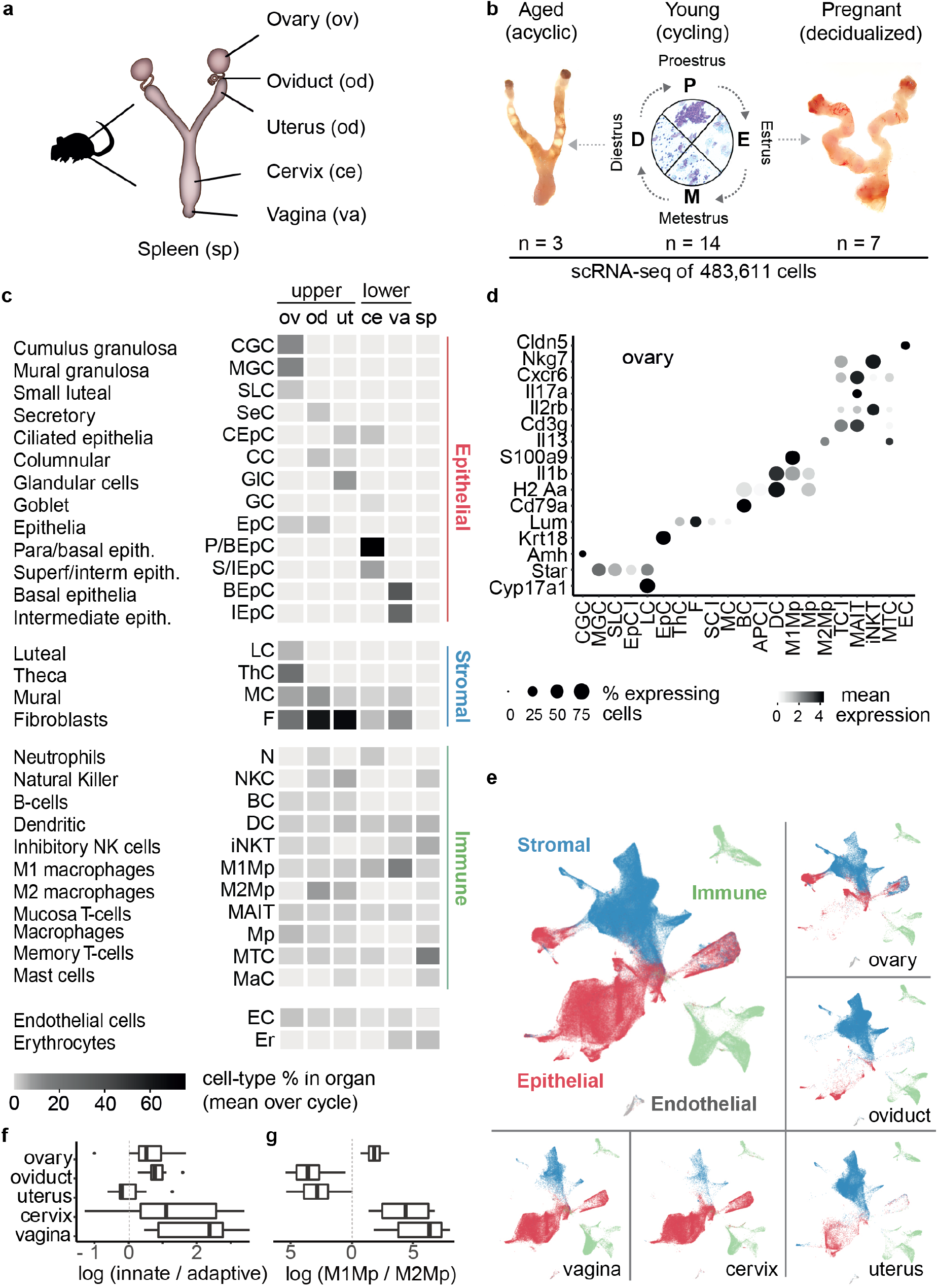
Single-cell characterization of the female reproductive tract. **(a)** Single-cell analysis of the reproductive tract (FRT) was performed on ovary, oviduct, uterus, cervix and vagina, with spleen as a non reproductive, control organ. **(b)** All organs of the FRT were profiled in multiple biological replicates at the four phases of the mouse estrus cycle, during aging, and during decidualization in pregnancy. Occurrence of leukocytes, and nucleated and cornified epithelial cells (see Figure S3a) was used to stage tissues in Proestrus (P), Estrus (E), Metestrus (M) and Diestrus (D). **(c)** Proportional heatmap of the most abundant cell types by organ (full list in Figure S1c), revealed a pronounced shift between the upper and lower FRT from stromal dominance to epithelial dominance in young cycling mice. **(d)** Selected marker genes used to classify ovarian cell types. Complete plots of marker genes used to classify all FRT organs are shown in Figure S2. **(e)** UMAP plot of the young cycling mouse cells. Cell types were integrated between organs for visualization purposes only. Cell types were assigned to epithelial (red), immune (green), stroma (blue), and endothelial (gray) compartments. **(f)** Log ratio of innate (N, DC, M1Mp, M2Mp, Mp and MaC) and adaptive (NKC, BC, iNkT, MAIT, MTC) immune cells abundances in the FRT organs. **(g)** Log ratio of M1 to M2 macrophages in the FRT organs.

We analyzed the single-cell transcriptomes from all cycle phases and organs (76,600 cells per organ on average) in young cycling mice to identify cell types and their organ-specificity. To ensure the accuracy of the cell type annotation, we combined two automated approaches with an extensive manual comparison of marker gene expression (Methods). These approaches identified approximately fifty cell types, including all expected stromal, epithelial and immune-cell sub-populations (Figure 1c,d, S1a-c, S2, Table S2). Cell types were defined independently in each organ and all organs were subsequently integrated together for visualization purposes (Figure 1e).

By evaluating the cellular composition of all five FRT organs simultaneously, we identified a collection of cell types shared across the entire FRT, and quantified the extent to which the FRT shows a pronounced shift in its cellular composition between the upper and lower tracts (Figure 1c,e, S1c). The cell types shared across the FRT include stromal fibroblasts, dendritic cells, macrophages, mural cells, a number of T cell subtypes, and endothelial cells, among others.

As expected, substantial variation in cellular composition occurs among the FRT organs. For instance, stromal cells outnumber epithelial cells in the upper FRT, whereas epithelial cells are more numerous in the lower reproductive tract (Figure 1c,e, S1c,d). The composition of the immune compartment also profoundly shifts between the upper and lower tracts. The upper tract is characterized by an enrichment of adaptive immune cells, including mucosa-associated T cells, memory T cells, and Natural Killer cells. The lower reproductive tract is dominated by the innate immune system, with a higher proportion of dendritic cells and neutrophils (Figure 1f, S1c). In addition, the balance between M1 and M2 macrophages profoundly changes between organs (Figure 1g, S1c). In the uterus and oviduct, wound-healing associated M2 macrophages dominate the cellular landscape, where they are likely involved in hormonally induced tissue remodeling (Madsen et al., 2013). In contrast, the classically activated and pro-inflammatory M1 macrophages dominate the ovary, consistent with previous reports that M1 but not M2 macrophages are required for folliculogenesis (Ono et al., 2018). M1 macrophages are also prevalent in cervix and vagina, consistent with these organs’ higher exposure to pathogens. Natural killer cells are concentrated in the uterus (and to a small extent in the oviduct), where they have been suggested to regulate decidualization (Sojka et al., 2019).

Our analyses identified major cell types present across the entire FRT, including stromal fibroblasts and macrophages; and revealed an anti-vs pro-inflammatory transition between the upper and the lower reproductive tract.

### Estrus cycling dramatically remodels the cervix and vagina immune compartments

To understand how tissue-environment, hormone state, and intrinsic cell identity combine to shape each organ’s tissue landscape, we compared how the cellular composition of the entire female reproductive tract varies across the cycle in young female mice. We precisely staged the cycle by the emergence of specific cell types in vaginal smears (Figure S3a, Methods). The correct identification of the phases was confirmed by an unbiased reconstruction of a transcriptional pseudo-time trajectory of uterine fibroblasts, which ordered the mice according to the smear-assigned phases (Figure S3b). Indeed, the substantial changes observed in vagina by single-cell data (Figure 2a) are consistent with the vaginal smears used to stage each mouse’s cycle. For example, the proportion of nucleated epithelial cells (such as BEpC and IEpcC) is at its highest in proestrus and lowest at metestrus (statistically significant by compositional regression, Methods, Table S3). We also observed other expected features, including a large expansion of the stroma from proestrus to estrus (Jin, 2019; Wood et al., 2007) and an expansion of glandular cells at proestrus in the uterus (Sato et al., 1997) (Figure S3c).

**Figure 2.**
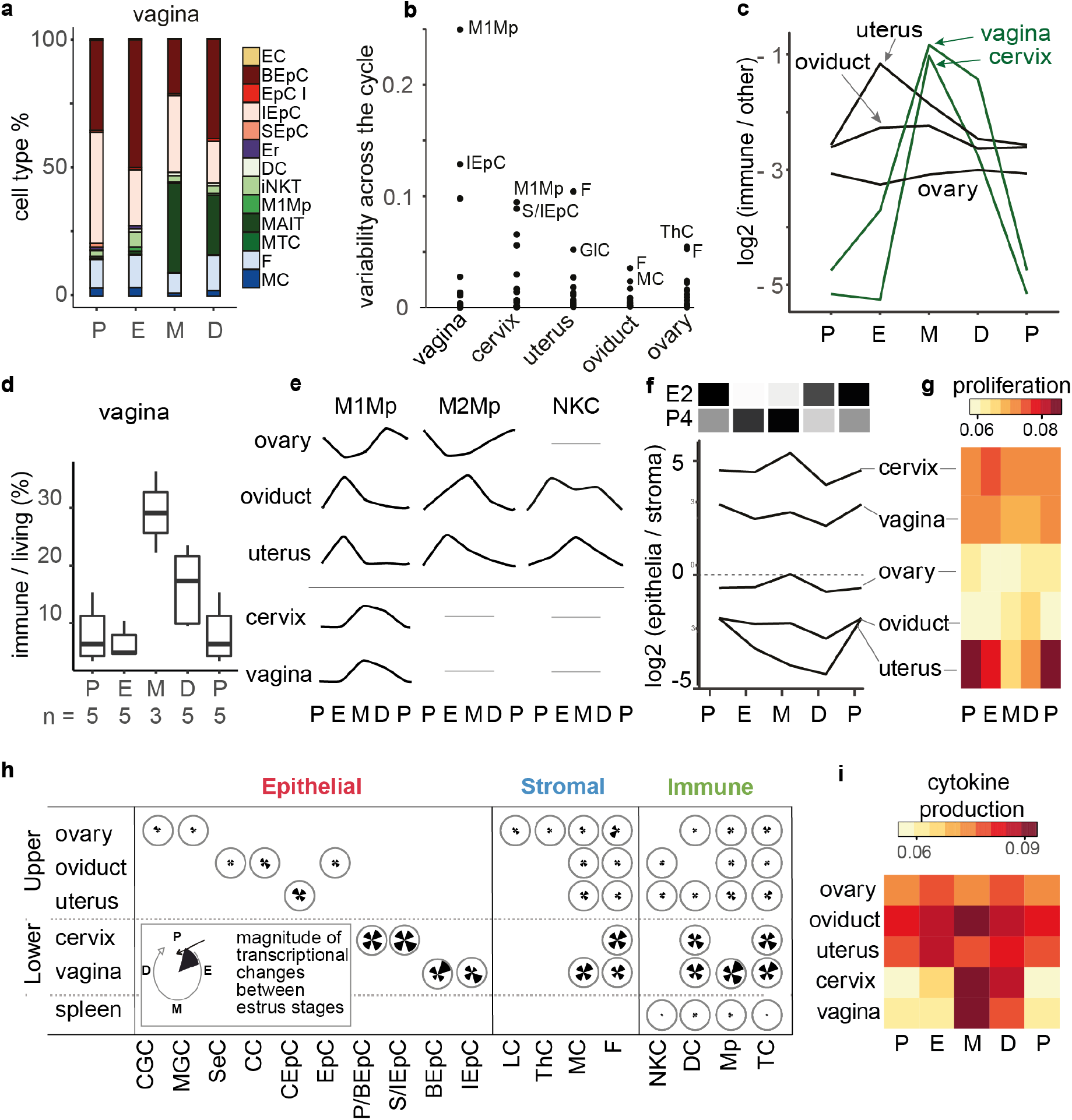
Estrus cycle drives organ-specific compositional changes. **(a)** Barplot showing the % of each cell type in vagina at each phase of the cycle (P-proestrus, E-estrus, M-metestrus, D-diestrus). The values shown are the averages across biological replicates, barplots per biological replicate are shown in Figure S3c. **(b)** The compositional variability across the cycle was plotted for each cell type in each organ. The y-axis shows the variability plotted as an interquartile range; for each organ, the two most variable cell types are indicated. **(c)** The ratio of immune to other cells across the cycle was plotted. The values shown are average across biological replicates, standard errors are shown in Figure S4c. The immune compartment is invariant in the ovary and oviduct, peaks during estrus in the uterus, and during metestrus in vagina and cervix. The ratios shown are the average across biological replicates, standard errors are omitted for clarity and shown in Figure S4a. **(d)** The cyclical changes in the vaginal immune compartment were independently confirmed using flow cytometry (‘n’ indicates the number of biological replicates). **(e)** Compositional changes across the cycle in all FRT organs were plotted for M1Mp, M2Mp and NK cells. **(f)** The ratio of epithelia to stroma across the cycle was plotted; uterus uniquely and extensively reshapes its cellular composition. The relative concentration of estradiol (E2) and progesterone (P4) at each stage from (Nilsson et al., 2015) is shown above (black is the maximum value of the cycle, white is 0). **(g)** Average activity score of genes promoting cell proliferation (GO:0008284) calculated in epithelial cells using AUCell. **(h)** Similarity of gene expression between the cycle phases for each cell type was quantified using optimal transport, and displayed as a flower plot (inset). Petal lengths indicate magnitude of transcriptional changes; for example, most cell types are more transcriptionally dynamic in cervix and vagina than in the upper FRT. **(i)** Average activity score of cytokine regulatory genes (GO:0001816) calculated in immune cells using AUCell.

We analyzed the cellular population dynamics from both an organ-specific and cell type specific point of view. To quantify the magnitude of remodeling in each organ, we determined the fraction of each cell type at each phase of the cycle and plotted the cycle inter-quartile range. This comparative approach revealed that vagina has the most cycle-variable cell types, as well as the highest average amplitude of compositional changes in the FRT (Figure 2b). The variability in cell composition across the cycle tends to decrease along the FRT: in uterus and cervix fewer cell types vary across the cycle, while oviduct and ovary are relatively invariant in their composition.

We then asked how the immune compartment of the female reproductive tract is remodeled across the cycle (Figure 2c, S4a). We found that the fraction of immune cells is lowest during proestrus in all organs, and is also low during diestrus in the upper reproductive tract. The vagina and cervix show considerably greater variation in their content of immune cells, with a maximum at metestrus. For vagina, we independently confirmed the sharp peak in immune cell numbers in metestrus by quantifying the fraction of immune cells across the cycle via flow cytometry (Figure 2d, S4b). In the uterus, in contrast, the immune-cell proportion peaks earlier at estrus, followed by a slower decline back to the minimum at proestrus. Interestingly, the immune fraction of the ovary and oviduct remains relatively invariant. Within specific immune cell populations, immune cells in the oviduct transition from M1 during estrus to M2 during metestrus (Figure 2e, S3c). In the uterus, M1 and M2 abundance also peaks during estrus, whereas metestrus is characterized by a specific abundance of NK cells.

Our analyses newly quantified how the different organs of the FRT are immunologically distinct. The lower reproductive tract undergoes cyclical, acute immune influx, which is also seen to a smaller but significant extent in the uterus and oviduct, while the ovary maintains an invariant population of immune cells throughout the cycle. Our characterization of the cycle dynamics of cellular abundance can be explored in an interactive online tool (see Data Availability).

### Uterus undergoes profound cyclical epithelial/stromal remodeling

Cell proliferation and death can be regulated by systemic steroid hormones (Wood et al., 2007). Leveraging on our simultaneous profiling of the five organs of the FRT, we asked whether tissue proliferation/remodeling is synchronized between them. To determine if the relative abundances of stroma and epithelia are coordinated across organs and across the cycle we calculated the ratio of epithelium to stroma across the FRT (Figure 2f, S4c). In all FRT organs, this ratio is lowest during diestrus and increases on the transition to proestrus; this coincides with the known progesterone minimum at diestrus (Figure 2f) (Nilsson et al., 2015). With the exception of uterus, FRT organs are relatively stable in the transition from proestrus to estrus and peak at metestrus. In uterus, proestrus profoundly changes the ratio of epithelia to stroma, and therefore we asked if proliferation contributes to these dynamic changes. We derived proliferation scores using the scRNA-seq data (Methods), which revealed that proliferation rates of epithelia are highest during the estrogen (E2) surge at proestrus and lowest at metestrus (Figure 2g). In contrast, stromal proliferation peaks at estrus and metestrus (Figure S4d), coinciding with peak progesterone (P4) levels, which promotes stromal proliferation and inhibition of the E2-induced epithelial proliferation (Li et al., 2011).

The most notable contrast in tissue remodeling is between vagina and uterus, providing a high-resolution quantification for the current models obtained from histopathological analyses (Sato et al., 1997). In vagina, there are cell type specific changes within the epithelial compartment across the cycle (Figure 2b) which do not impact the overall balance of epithelia to stroma (Figure 2f). In contrast, the uterus shows strong changes in its epithelial to stromal ratio (Figure 2f).

In sum, our data reveals how proliferation and immune activation are precisely regulated during the cycle. The regulation of cell type abundances and immune infiltration in the FRT is highly organ specific, despite their equal exposure to circulating hormones.

### Tissue morphological changes are tracked by gene expression

We considered the possibility that organs such as ovary and oviduct, where cell abundances are relatively stable across the cycle, might instead show substantial changes in transcription. We scored the magnitude of transcriptional change between phases of the cycle for each cell type in every FRT organ using optimal transport analysis (Methods) (Figure 2h). This analysis revealed that cycle– related variation in cell type abundance is accompanied by a corresponding scale of transcriptional change. Vagina and cervix are high in both cell type variability and gene expression; in contrast, the cells in uterus, oviduct, and ovary show less of both (Figure 2b,h). When considered by cell types, we observed the same effect: the abundance of immune cells closely corresponds to their functional activation, which we orthogonally measured by single-cell scoring of inflammation markers and cytokine gene expression (Figure 2i, Table S9), in agreement with total cytokine and chemokine concentration found in vaginal and uterine lavages (Hickey et al., 2013; Sonoda, Y. et al., 1998).

Our data show that organs with the largest morphological changes also have the largest transcriptional changes across all cell types, suggesting a positive feedback loop interaction between morphological and transcriptional change.

### Fibroblast functions are dynamically regulated throughout the cycle, but not coordinated between organs

Fibroblasts are key regulators of wound healing and inflammation in multiple organs, we therefore asked if fibroblasts play the same functional roles in the cycling FRT organs, and whether these roles are coordinated between them. We first used over-representation analysis (ORA) to identify the functional pathways enriched among cycling genes with significant differential expression between any two adjacent phases of the cycle (Figure 3a, Table S7, Methods). This revealed that ECM remodeling and Tumor necrosis factor (Tnf) regulation of inflammation are core programmes of fibroblasts during the cycle. However, the activity of these two specific functional pathways is often out of phase or even anti-correlated between organs, despite equal exposure of the organs to circulating hormones (Figure 3b,c).

**Figure 3.**
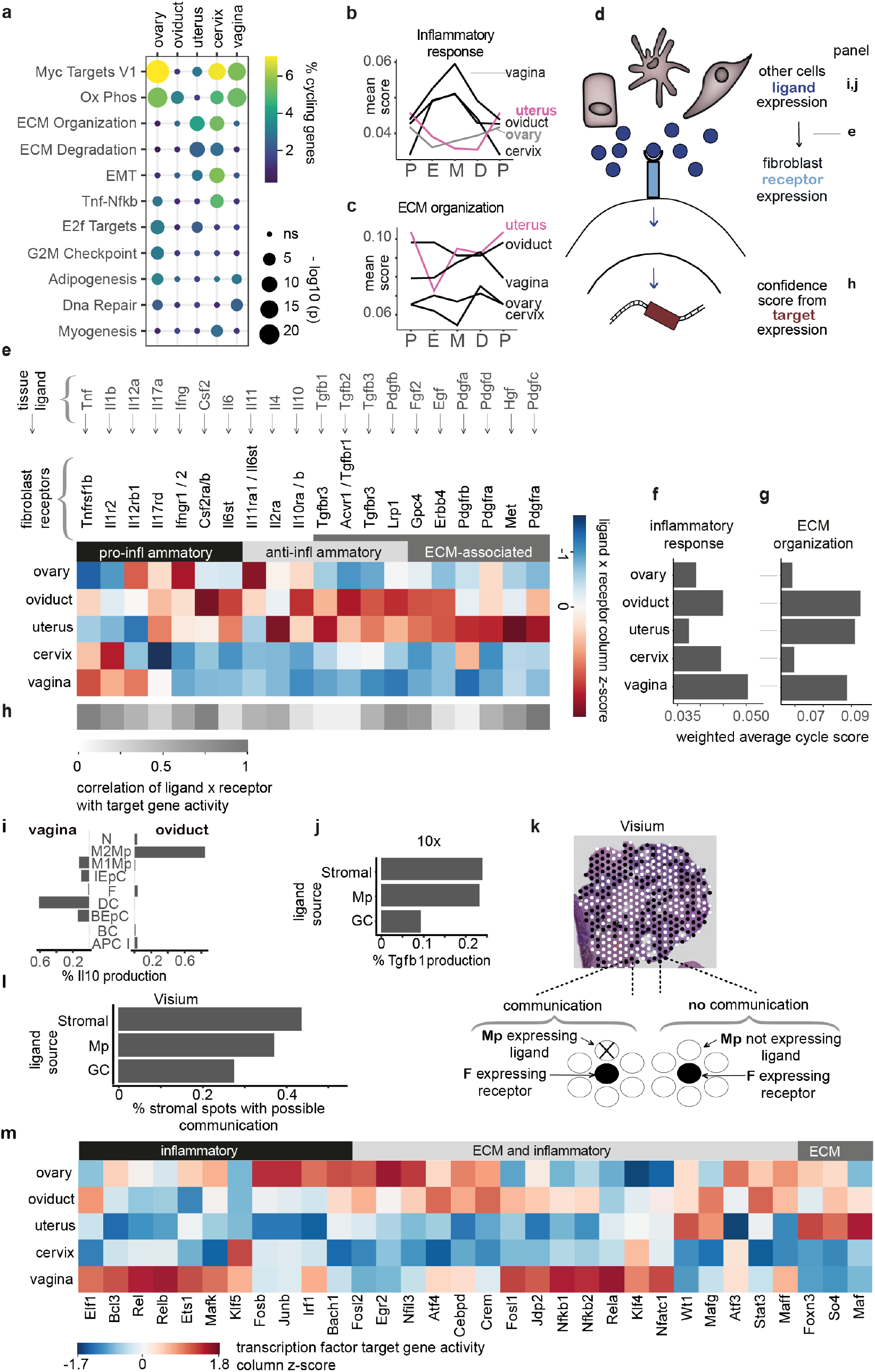
Gene expression dynamics of fibroblasts across the female reproductive tract. **(a)** Functional pathways, including inflammation and ECM, enriched in genes differentially expressed between phases of the cycle in fibroblasts. **(b)** Activity scores of inflammatory genes (Sup. Table S9) determined by AUCell and averaged across all fibroblasts in each cycle phase (P-proestrus, E-estrus, M-metestrus, D-diestrus). **(c)** Activity scores of ECM organization genes (Sup. Table S9), as in (b). **(d)** Schematic of the cell-to-cell ligand-receptor and ligand target analyses. **(e)** Heatmap showing the z-scores of ligand-receptor products averaged across phases. Ligand expression is averaged across all cell types; receptors are in fibroblasts only. A full version of the heatmap is shown in Figure S5a. The receptor-ligand interactions shown were deemed statistically significantly different in at least one FRT organ in comparison to the rest by a permutation test (p-values in Table S4). **(f)** Activity scores of inflammatory genes averaged across all fibroblasts in each phase in each organ. The scores shown here are the average of the scores shown in panel (b), weighted by phase length (Methods). **(g)** Activity scores of ECM genes averaged across all fibroblasts in each phase in each organ. The scores shown here are the average of the scores shown in panel (c), weighted by phase length (Methods). **(h)** Heatmap of Spearman correlation values between the expression product of a ligand-receptor pair and the AUCell activity score of predicted targets of the ligand (Browaeys et al., 2020). **(i)** Top 5 cell types with the largest expression of *Il10* in vagina and oviduct. **(j)** Contribution of stromal cells, macrophages and granulosa cells to Tgfb1 expression calculated using 10x scRNA-seq data. **(k)** Schematic showing cell-to-cell communication scoring strategy in spatial transcriptomics analysis. Communication between spots containing fibroblast and ligand source cells was considered possible if: the fibroblast spot expressed receptor (*Tgfbr3* or *Tgfbr1*/*Tgfbr2*), and neighborhood spots expressed ligand (*Tgfb1*) together with the correct cell type marker. **(l)** Proportion of stromal spots which are communicating with respective ligand source spots. Spots were scored using the strategy shown in (k). **(m)** Estimated activity scores of targets of transcription factors associated with inflammation and ECM regulation in fibroblasts. Shown are TFs whose activity scores were statistically significantly different between the FRT organs through a permutation test (Table S5, Methods).

The activation of inflammatory pathways in fibroblasts shows two organ-specific patterns (Figure 3b, Table S9, Methods): in vagina, cervix, and oviduct, inflammatory activity peaks at metestrus, and in uterus and ovary at proestrus. Thus, the vaginal immune cell infiltration we previously observed at metestrus (Figure 2c) is accompanied by fibroblast inflammatory activation. In contrast, ECM organization shows no coordinated pattern across FRT organs (Figure 3c). Compared to the other organs, uterus has the most extensive ECM remodeling which peaks at proestrus and reaches its minimum during estrus.

Our data confirms that cycling fibroblast programmes are similar between organs, yet often out of phase, suggesting that fibroblast functions are regulated by a combination of systemic and local cues.

### Cell-to-cell communication and transcription factor activity reveal high inflammation in the lower FRT and extensive ECM remodeling in the uterus

Fibroblasts coordinate organ function and homeostasis via communication with other cell types through ligand– receptor interactions (Davidson et al., 2021; DeLeon-Pennell et al., 2020). We therefore performed cell-to-cell communication analysis to identify the organ-specificity and activity of ligand-receptor interactions (Figure 3d, Methods) (Jin et al., 2021; Shao et al., 2021). We first focused on how ligands from all cell types converged on fibroblasts, by calculating communication scores as the product between a and b, where a is the expression of ligands averaged over all cell types in all phases (the ‘ambient’ ligand expression), and b is the expression of receptors averaged over fibroblasts in all phases.

Vaginal and cervical fibroblasts have the highest communication scores for pro-inflammatory Interleukin 1 beta (*Il1b*) and *Tnf*; and the lowest scores for anti-inflammatory Interleukin 10 (*Il10*), Interleukin 11 (*Il11*) and Transforming growth factor beta (*Tgfb*) cell-to-cell signaling (Figure 3e, S5a, Table S4, Methods). In contrast, uterine fibroblasts receive primarily anti-inflammatory signaling, and oviduct and ovary have a mixture of anti- and pro-inflammatory signaling. By summarizing the cycle average fibroblast inflammation scores by organ, we found that vagina has the strongest cyclical inflammatory transcriptional responses (Figure 3f), consistent with its cell type compositional changes (Figure 2c). A similar summative analysis of ECM reorganization demonstrated that uterus and oviduct undergo the largest structural remodeling across the cycle, followed by vagina (Figure 3g). Indeed, fibroblasts in the uterus and oviduct show the highest communication scores for ECM-associated signaling (Figure 3e, S5a).

To verify that increased cell-to-cell communication results in upregulation of downstream pathways, for each of the ligands, we scored the activity in fibroblasts in each FRT organ for their predicted target genes (Browaeys et al., 2020). Most ligand targets have a strong positive correlation with their organ receptor-ligand scores (Figure 3h, Methods). In other words, if an organ has high ligand-receptor activity, then it also has high ligand-target activity. We then sought to identify which cell types were responsible for signaling to fibroblasts by partitioning the transcription of each ligand by cell-of-origin (Figure S5b, S6a, Methods). In the lower reproductive tract, M1 macrophages (source of *Il1b, Tnf, Il12a*) and memory T-cells (source of *Ifng, Csf2*) appear responsible for most pro-inflammatory signaling. In the upper reproductive tract, M2 macrophages (source of *Il10*) and fibroblasts/theca cells (source of *Il11*) generate the predominantly anti-inflammatory environment (Figure S5b, S6a).

Our data reveal that the cell types responsible for inflammatory ligand production are often organ-specific. Prior studies in humans and cows have suggested organ-specificity in TNF signaling arising from uterine glandular cells (Okuda et al., 2010; Tabibzadeh, 1991, 1999), which we confirmed in the mouse (Figure S5b). As another example of organ-specific signaling, we identified the cell-of-origin of *Il10* (Figure 3i). *Il10* is highly active in oviduct (see Figure 3e), where the major source of ligand are M2 macrophages (Figure 3i, S5b). In vagina, where there is substantially less IL10 signaling and few M2 macrophages, the strongest source is dendritic cells (Figure 3i, S6a). In contrast to inflammatory signaling which is often organ-specific and paracrine, fibroblast ECM is often autocrine controlled by signaling from stromal cells (Figure S5b, S6a).

TGFB is one of the most potent regulators of ECM activity and inflammation (Derynck and Zhang, 2003). Using scRNA-seq data we determined that Tgfb1 is highly active in the organs of the upper FRT (Figure S5a). In the ovary, the main sources of *Tgfb1* are stromal cells and macrophages, and to a smaller extent granulosa cells (Figure 3j). To validate the TGFB signaling in the ovary between fibroblasts and their partner cells we generated spatially resolved transcriptomics data using Visium. Using spatial and transcriptional information, we calculated the proportion of spots with stromal cells that expressed Tgfb1 receptors and neighboured a spot with stromal cells, macrophages or granulosa cells that expressed Tgfb1 ligand (Figure 3k). We confirmed that indeed ligand-expressing stromal cells and macrophages co-localize significantly more frequently with receptor-expressing fibroblasts compared to granulosa cells (Figure 3l).

In addition to the cycle-averaged communication above, we evaluated the dynamics of cell-to-cell communication across the cycle in uterus and vagina, which had the highest levels of ECM and inflammation, respectively. In uterus, we observed that ECM-related cell-to-cell communication is lowest in the estrus phase (Figure S6b), consistent with a corresponding decline in ECM-related gene activity (Figure 3c). Similarly, in vagina, the highest pro-inflammatory and lowest anti-inflammatory cell-to-cell communication are found in metestrus (Figure S6b), consistent with inflammation peaking in metestrus (Figure 3b).

Finally, to identify candidate regulators involved in organ-specific inflammatory and ECM processes, we quantified the associated transcription factor (TF) activity across the cycle using SCENIC (Figure 3m, S6c, S7a, Table S5). As expected, we found that the activity of inflammation-associated transcription factors is highest in vagina, while activity of ECM-related transcription factors is highest in the uterus (Figure 3m). For example, in vaginal fibroblasts, both canonical (*Nfkb1, Rela*) and noncanonical NFKB pathways (*Nfkb2, Relb, Bcl3*) are activated by IL1B and TNF ligands (Figure S6c), suggesting that the synergistic action of these ligands on NFKB pathways contributes to inflammation (Di Paolo et al., 2015). Overall, we observed that the dynamics of transcription factor activity correspond closely with changes in cell-to-cell communication (Figure S7b).

We confirmed that fibroblasts are central regulators of inflammation and ECM across the FRT. We discovered that the timing and underlying transcriptional regulators of fibroblast activation are highly organ specific. The differentially expressed genes, co-expressed gene clusters, cell-to-cell communication, and transcription factor activity can be explored in our interactive online tool (Data Availability).

### Modeling the human menstrual cycle using mouse decidualization

The human reproductive cycle includes a step of terminal differentiation (decidualization) of the uterine stromal cells, which in other mammals only happens in the presence of a fertilized egg. During the first trimester of human pregnancy, the immune microenvironment of the decidua prevents inflammatory responses (Vento-Tormo et al., 2018). Here we first asked whether mouse decidual cells also display an anti-inflammatory profile, and then sought to quantify the degree of transcriptional conservation between humans and mouse cycling fibroblasts.

To parallel the spontaneous decidual reaction that occurs during the human cycle, we induced decidualization in mice by inducing pregnancy, and characterized the uterine architecture at embryonic day 5.5 by scRNA-seq in seven biological replicates (Figure 4a, S8a,b). A subset of stromal cells unique to pregnant mice transcriptionally expresses the classical markers of decidualization: *Alpl, Bmp2* and *Prl8a2* (Figure 4b, S8c) (Finn and Hinchliffe, 1964; Ramathal et al., 2010; Soares et al., 1998). As expected, many stromal cells from pregnant uteruses do not express these markers, because the mouse uterus decidualizes heterogeneously (Zhao et al., 2017). For instance, *Bmp2* is only expressed in stromal cells surrounding the implanted embryo. We confirmed decidualization histologically using H&E staining (Figure 4c). Compared to the metestrus phase, the pregnant uterus is characterized by the appearance of decidual cells, accompanied by a proportional increase in NK, glandular cells (GlC) and fibroblasts, and a decrease in columnar (CC), ciliated epithelial cells (CEpC), mural (MC), dendritic (DC) and MAIT cells (Figure 4d). Additionally to cell compositional changes, decidualization in mouse causes extensive transcriptional changes in inflammation, ECM and embryo development pathways (Figure S8d).

**Figure 4.**
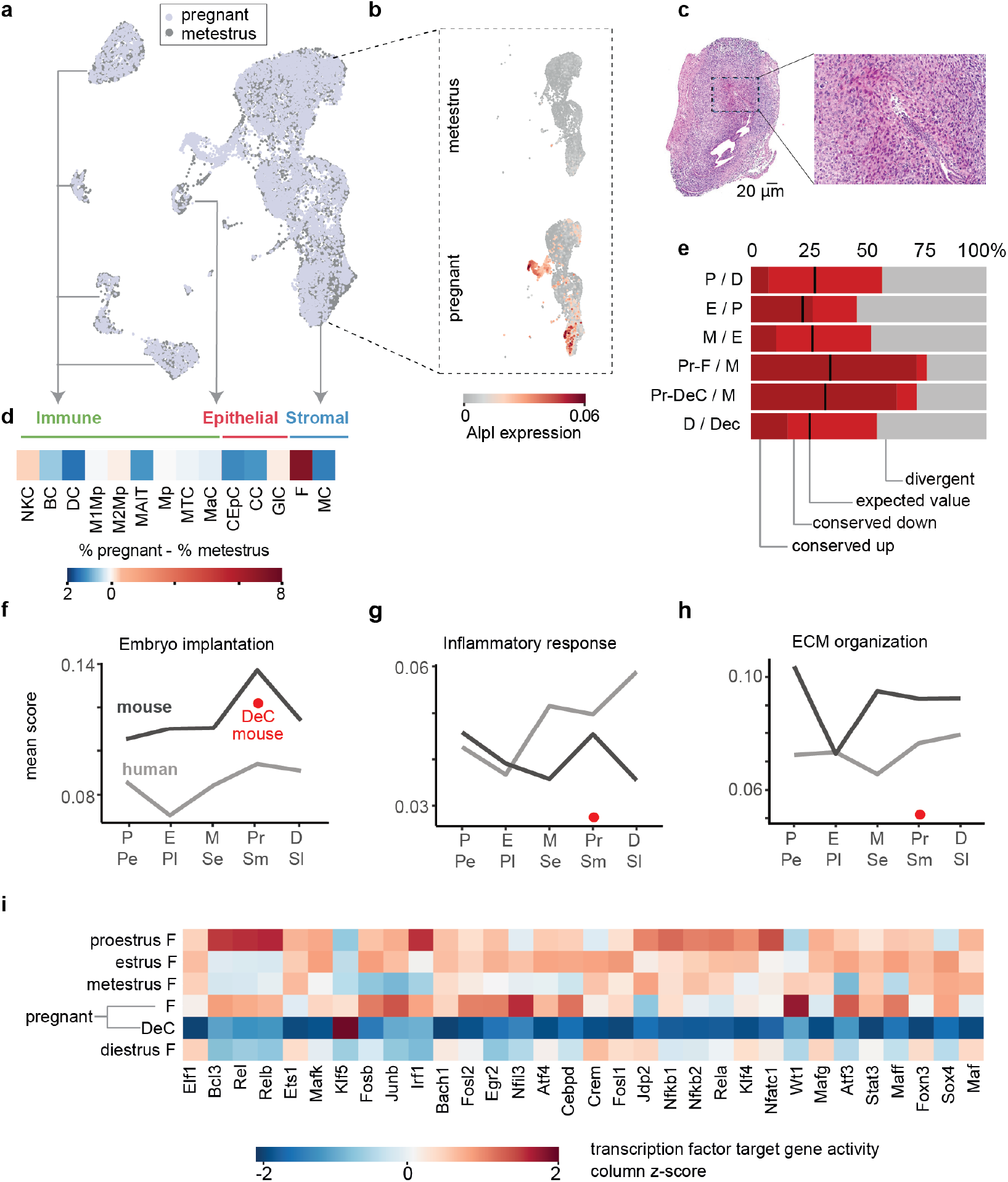
Regulation of reproductive cycle and decidualization is conserved between mouse and human. **(a)** UMAP plot of the integration of the pregnant with the metestrus samples. Shown in this panel is a subsample of 19,724 of the total 40,828 cells used in the analyses. **(b)** UMAP from panel a) subsetted to stromal cells and split by condition showing the expression of marker gene of decidualization *Alpl*. **(c)** H&E staining of pregnant mouse uterus showing decidualization at the implantation site. **(d)** Cell abundance compositional changes in early pregnancy compared to metestrus. Heatmap shows the difference in average % of each cell type between pregnant and metestrus samples. Pregnancy in mice is characterized by the appearance of decidual cells. To be able to compare the changes in all other cell types upon decidualization, decidual cells were omitted from the comparison and cell abundances were re-calculated. **(e)** Conserved differentially regulated genes in mouse uterine fibroblasts and decidual cells and human fibroblasts across the cycles. Barplot indicates in red the % of homologous differentially regulated genes that showed the same or opposite directionality of regulation in comparison to adjacent cycle phases in paired mouse-human cycle phases. For instance, a mouse gene upregulated in proestrus compared to diestrus and the homologous human gene upregulated in proliferative early compared to secretory late. Genes that showed opposite directionality of regulation (e.g. up-regulation in humans, down-regulation in mice) were considered divergent and their % is shown in gray. Black line shows the proportion of the conserved genes expected by chance in each cycle phase. As bar labels only mouse phase comparisons are shown (P-proestrus, E-estrus, M-metestrus, D-diestrus, Pr-pregnant, F-fibroblast, DeC-stromal decidual cells). **(f)** Activity scores of genes that regulate embryo implantation (GO:0007566) (Sup. Table S9) determined by AUCell and averaged across all mouse and human fibroblasts in each phase of the cycle (P-proestrus, E-estrus, M-metestrus, D-diestrus, Pe-proliferative early, Pl-proliferative late, Se-secretory early, Sm-secretory mid, Sl-secretory late). Red dot indicates the activity score in mouse decidual cells. **(g)** Activity scores of inflammation genes as in (f). **(h)** Activity scores of ECM genes as in (f). **(i)** Estimated activity scores of targets of transcription factors associated with inflammation and ECM regulation in mouse fibroblasts, across the cycle and in pregnancy, as well as decidual cells in pregnancy.

We asked whether the decidualized cells and fibroblasts in mice express the same transcriptional programs previously identified in human uterine fibroblasts (Figure 4e, Figure S9a, Table S6) (Wang et al., 2020). First, we used the same mutual information approach as the original study to re-identify 1670 human genes that are differentially expressed between specific phases of the menstrual cycle. We then tested whether the same genes are differentially regulated in the corresponding estrus phases in mice using the same approach (Figure S8e, Methods). At every phase of the cycle, the dynamic gene expression changes in human and mouse are more conserved than expected by chance (Methods), and the phase of decidualization has an especially high percentage of conserved transcription (Figure 4e). These genes are enriched for ECM, inflammation, and cycle regulation and implantation pathways (Figure S9b). Most of these processes show species-specific differences in activity across the cycle; however, the transition to decidualization is largely conserved (Figure 4f-h, Figure S9c). When compared to fibroblasts, decidual stromal cells show consistently lower activity of ECM- and inflammation-related genes (Figure 4g,h) and transcription factors (Figure 4i).

In sum, our analyses revealed that mouse decidual cells display a markedly anti-inflammatory transcriptional profile and that the transition to decidualization is largely conserved between mouse and human.

### Inflammaging in the FRT

The FRT shows signs of accelerated aging compared to other organs. It undergoes extensive physiological changes following its decline in mid-life, which culminates in menopause/acyclicity (Broekmans et al., 2009). We therefore asked which age-related changes occur in the aged FRT compared to young. Our pathological analysis confirmed that 18 month old mice often display immune infiltration in most organs (Finch, 2014; Nelson et al., 1984), ovarian atrophy, as well as localized ovarian and/or uterine hyperplasias (Figure 5a, Figure S10, Table S10). To quantify how age-associated cessation of the estrus cycle changes the cellular composition and transcription in reproductive organs, we collected in triplicate and analyzed the single-cell transcriptomes of all reproductive organs and spleen from 18-month old female mice (Figure 5b, S11a).

**Figure 5.**
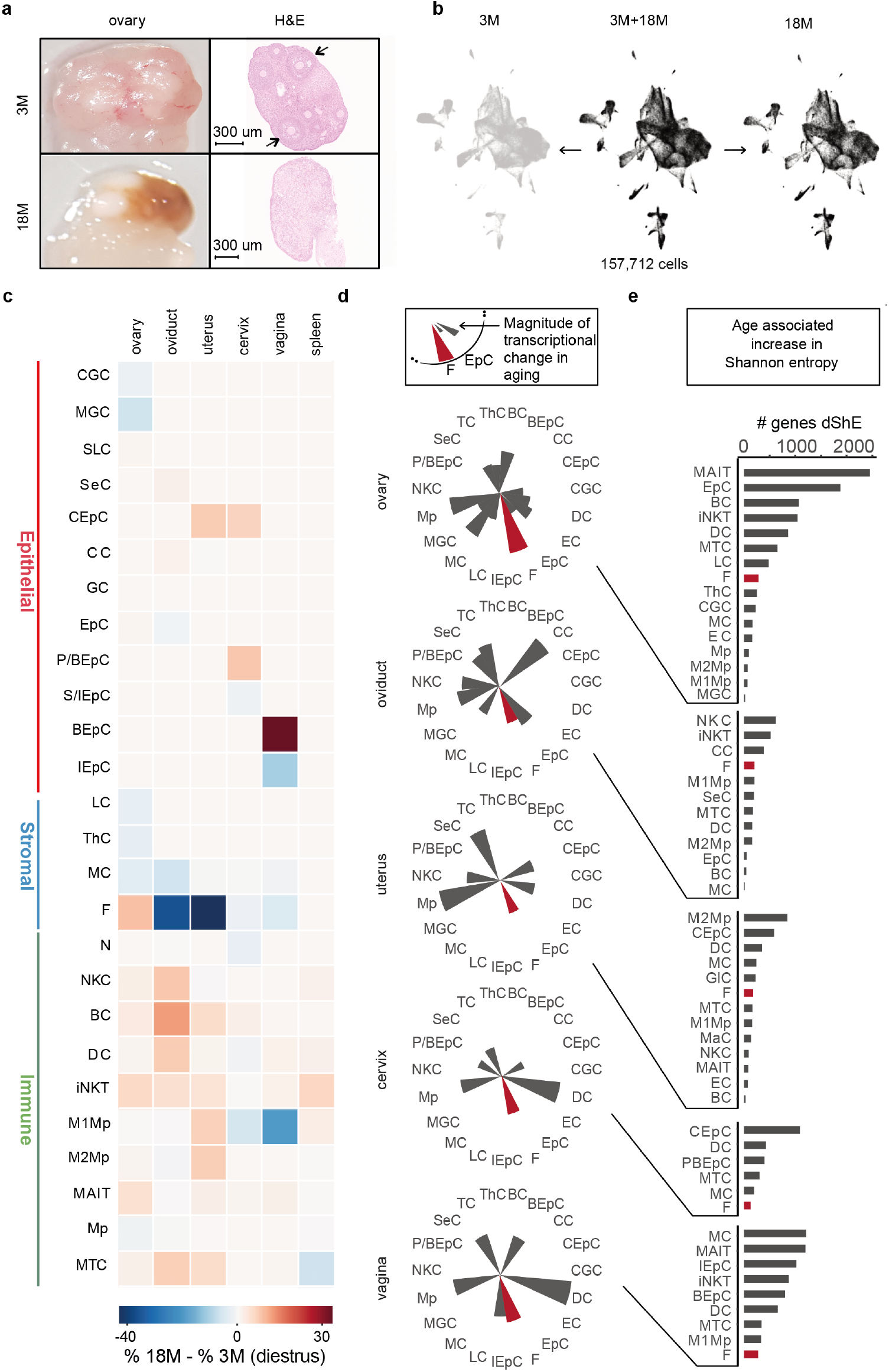
Organ-specific impact of aging on the FRT. **(a)** Photograph and H&E staining of young and aged ovaries. Arrows point to follicles in the young ovary. Aged ovary is atrophied and contains no follicles. **(b)** UMAP plot of the integration of the aged mouse samples with the diestrus samples. **(c)** Cell abundance compositional changes in aging compared to diestrus. Heatmap shows difference in average % of each cell type between aged and diestrus samples. **(d)** Similarity of gene expression programs between the aging and diestrus for each cell type quantified using optimal transport. Line lengths indicate magnitude of transcriptional changes. Optimal transport distances of fibroblasts are colored in red. **(e)** Number of genes with increased differential Shannon entropy (ShE) of all cell types in ovary, oviduct, uterus, cervix and vagina in diestrus compared to old age.

We identified extensive changes in cell type abundances by comparing cell type proportions between the aged and young mice in diestrus (Figure 5c, S11b), the phase most similar to acyclicity (Felicio et al., 1984). Ovary shows a decrease in the proportion of follicle-associated cells such as ThC, MGC, and LC, which is to be expected due to the exhaustion of ovarian follicles and corpora lutea in acyclic mice (Lliberos et al., 2021). We independently confirmed this decrease by both histopathology and RNAscope (Figure S10, S11d,e). As expected, we found that the proportion of fibroblasts increases in ovary (Lliberos et al., 2021) and decreases in the oviduct and uterus (Craig and Jollie, 1985).

By evaluating the entire FRT, we newly quantified how aging increases the fraction of immune cells in the upper reproductive tract, whereas aging decreases the immune cells in the lower reproductive tract. Prior studies on single organs found similar results in isolation (Elmes et al., 2015; Rodriguez-Garcia et al., 2021; Yaakov et al., 2021). The remodeling of the immune compartment is organ-specific: uterus shows an increase in M1 and M2 macrophages, cervix and vagina have a decrease in M1 macrophages, while oviduct has an increase in NK, B and dendritic cells. The control organ, spleen, displays statistically significant differences only in iNKT and PC proportions (Figure 5c, S11b,c, Table S3), in agreement with previous reports (Kimmel et al., 2019).

### Age-related gene expression changes are organ-specific

We then compared the gene expression programs between young and old mice using optimal transport analysis. In contrast to the transcriptional changes associated with estrus, which are concentrated in the lower FRT, during aging both the upper and lower FRT show extensive gene expression changes (Figure 5d). We found that the magnitude of age-related changes in cell type transcription is highly organ-specific. For example, fibroblasts show transcriptional changes during aging in all organs, but in ovary they are the most strongly altered cell type.

Increase in cell-to-cell transcriptional variability has also been shown to be associated with aging (Enge et al., 2017; Martinez-jimenez et al., 2017), though this variability may be cell type specific (Kimmel et al., 2019). Simultaneously profiling all five FRT organs allowed us to investigate how gene-wise transcriptional variability changes during aging for over 50 different cell types, and whether cell types that are common between organs transcriptionally age in a similar manner (Figure 5e, S12, Table S8). Importantly, we scored the age-associated transcriptional variability against the natural cyclical variation found in these organs, using Shannon-entropy, a metric commonly used for quantifying diversity in ecology.

Taking ovary as an example, aging strongly increases the cell-to-cell variability in the majority of cell types, when compared with other FRT organs. This age-related transcriptional variability is often cell type specific: MAIT and EpC cells increase substantially with age, whereas M1 and M2 macrophages are largely unaffected (Figure 5e, S12). As a cell type shared among all FRT organs, fibroblasts show modest cell-to-cell variability between young and old mice. Other cell types found across the FRT such as MAIT cells are more variable (Figure 5e, S12). The transcriptional variability of many epithelial cells (BEpC, CEpC and IEpCs) is substantially changed with age in uterus, cervix and vagina; however, these age-related differences are smaller than the variation observed in the normal cycle (Figure S12).

In sum, aging results in substantial changes to the cell type composition of FRT organs, most notably immune infiltration in the upper reproductive tract, and the age-related gene expression changes for each cell type are organ-specific. Unlike the cyclic FRT, both upper and lower aged FRT show profound changes in their transcription programs. Comparison of FRT organs shows that the ovary is more affected by changes in the gene-wise cell-to-cell variability.

### Organ-specific impact of chronic inflammation during FRT aging

Fibroblasts can retain inflammatory memory (Kirk et al., 2021), and thus shape age-related changes to organ physiology and function. To test the extent and impact of inflammatory responses in aged fibroblasts, we evaluated the difference in inflammation scores between young diestrus and old acyclic FRT organs and then applied a linear mixed model (Figure 6a). Aging results in a significant increase in fibroblast inflammation in all organs except ovary. Interestingly, the rate of increase is significantly different between FRT organs (p-value < 0.05 of the organ-age interaction terms, Methods), with cervix and uterus displaying the most pronounced increase in inflammation.

**Figure 6.**
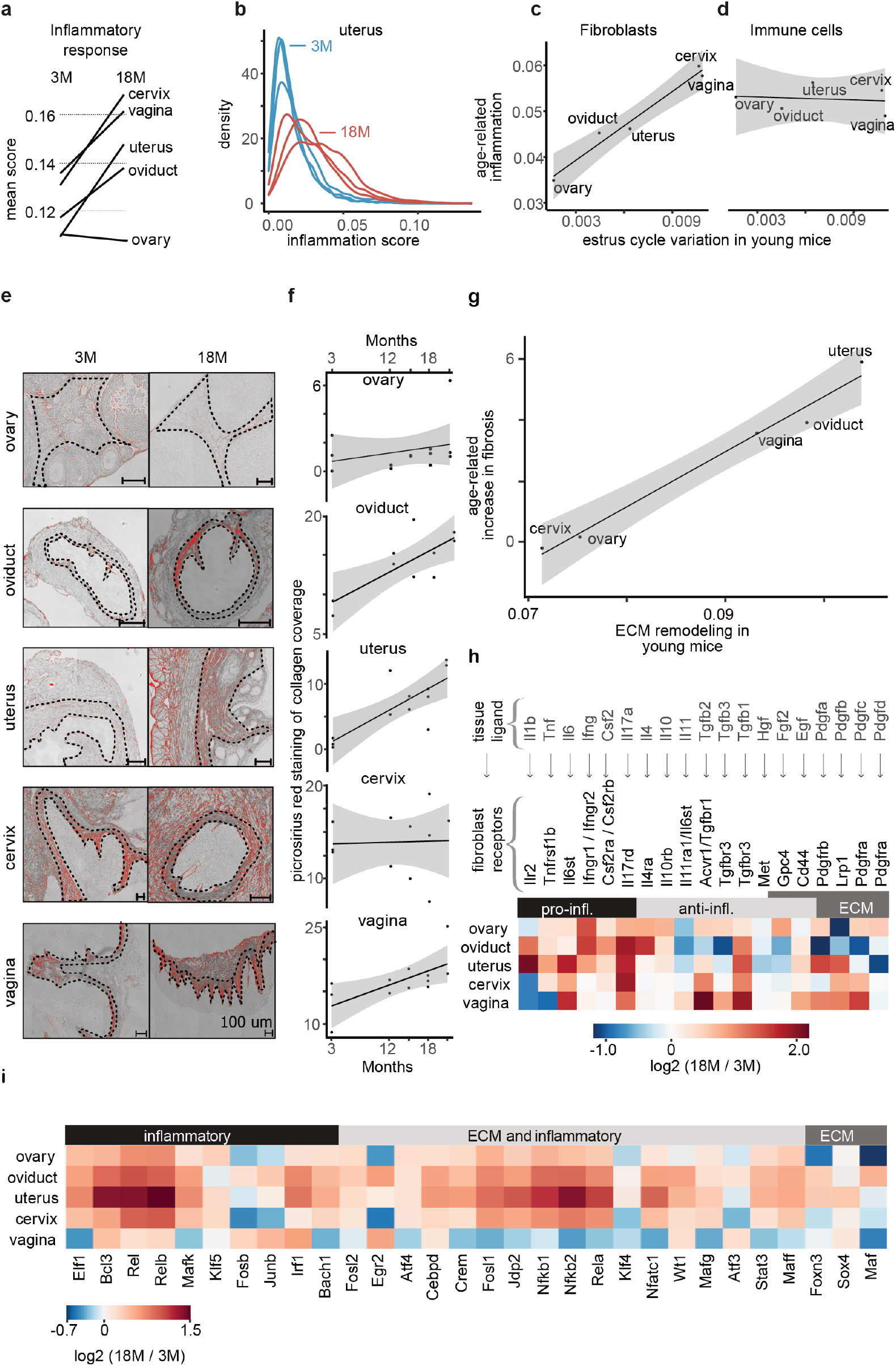
Inflammation and fibrosis in the FRT accumulate gradually with aging. **(a)** Activity scores of genes in inflammation determined by AUCell and averaged across all mouse fibroblasts in diestrus and old age (Table S9). **(b)** Density plot of fibroblasts inflammation scores in each biological replicate of young and old mice. **(c, d)** The relationship between the amplitude of organ-wide inflammation scores of young mice during the cycle (Figure 3b, Sup. Table S9) and the inflammation score of fibroblasts (Figure 6a, Table S9) and immune cells (Table S9) from the different organs in old age. **(e)** % area of stained collagen deposition in ovarian, oviductal, uterine, cervical and vaginal tissue of 3 and 18 month old mice. **(f)** Quantification of % area of stained collagen deposition in all FRT tissues of 3, 12, 15, 18 and 21 month-old mice. **(g)** The relationship between the maximum of organ-wide ECM reorganization scores of young mice during the cycle (Figure 3c, Sup. Table S9) and the fibrosis score of different FRT organs in old age. Fibrosis score was calculated as the difference in % area of stained collagen deposition in old (18 months) and young mice (3 months). **(h)** Heatmap showing the log2 fold changes of ECM-associated and inflammatory ligand-receptor products in old age compared to diestrus of all FRT organs. Ligand expression is averaged across all cell types; receptors are in fibroblasts only. **(i)** Activity scores of targets of ECM and inflammation-associated transcription factors in aged samples compared to diestrus.

We quantified what fraction of this inflammation is due to a subset of highly active fibroblasts versus a general increase in all fibroblasts. For each young and old uterus, we plotted the distribution of inflammation scores of the fibroblasts (Figure 6b). We statistically evaluated the observed differences using the waddR package (Schefzik et al., 2021), revealing that the distributions of inflammation scores are significantly different (p-value = 0.002) between young and old mice. We further dissected these distributions via decompositional analysis using a 2-wasserstein distance-based approach, revealing that their shape, location, and sizes equally contribute to their differences. In other words, the increase we observed in fibroblast inflammation is driven by both an expansion of sub-populations with high inflammation scores as well as an overall increase in expression of inflammatory genes throughout the population.

We tested the hypothesis that recurrent, cycle-related inflammation of fibroblasts across the reproductive lifespan might accumulate into age-related chronic inflammation. We quantified the amplitude of the organ-wide inflammation scores in young mouse fibroblasts during the cycle and compared it to age-related inflammation. Indeed, the higher an organ’s amplitude of inflammation is during the cycle at a young age, the higher the organ’s fibroblast inflammation score is in old age (Figure 6c). In contrast, immune cells do not display such an association (Figure 6d). These results suggest that fibroblasts incompletely resolve recurrent inflammation from the cycle, thereby retaining a cumulative memory of past inflammation.

### Tissue fibrosis accumulates gradually in oviduct, uterus and vagina

Inflammation is closely linked with fibrosis (Flavell et al., 2008; Kendall and Feghali-Bostwick, 2014), and increased collagen deposition and fibrosis can be the end result of chronic exposure of fibroblasts to inflammatory cytokines (Lliberos et al., 2021; Selman and Pardo, 2021).

We previously observed significant ECM tissue remodeling during the cycle of young mice (Figure 3c). We considered that - similar to inflammation - the incomplete resolution of this remodeling could lead to age-related pathological accumulation of collagen, and thus to fibrosis. We thus measured intercellular collagen in FRT organs using Picrosirius red staining to label collagen I and III fibers in 3 and 18 month-old mice, in triplicate, and in 12, 15 and 21 month-old mice in duplicate. Accumulation of fibrosis in uterus can lead to infertility (Sahin Ersoy et al., 2017; Secomandi et al., 2022), and we found that collagen increases in uterus both at an early stage and increases most steeply than the other organs over aging (increase of 3.2% per six months, p-value=0.013) (Figure 6e,f, S13). Oviduct (2.7%, p-value=0.022) and vagina (2.3%, p-value=0.033) also showed steady increases in collagen; ovary and cervix did not show collagen increases (Figure 6e,f, S13). Independent pathological analysis confirmed prevalent fibrosis in the stroma of oviduct and uterus (Table S10). For each FRT organ, the age-related collagen accumulation rate was best predicted by the maximum phase-specific ECM activity in the cycle (Figure 6g). In other words, the intensity of ECM remodeling in fibroblasts during the cycle corresponds with the severity of age-related fibrosis.

We asked whether our observed inflammation and fibrosis accumulation was reflected in organ-specific fibroblast regulatory and signaling networks. Using SCENIC, we found that the aging uterus showed a large increase in both the number and intensity of transcriptional regulatory modules associated with inflammation and ECM (Figure 6i). Other FRT organs showed more moderate (oviduct, cervix, ovary) or low (vagina) increases. Similarly, we found elevated inflammation and ECM activity in the ligand-receptor interactions centering on uterine fibroblasts (Figure 6h).

Our results indicate that ECM accumulation as a result of incompletely resolved cyclic ECM remodeling can lead to gradual fibrosis development. Intensity of ECM remodeling in each organ corresponds with the severity of fibrosis and predisposes each organ differently to fibrosis development.

## Discussion

In humans, like most other mammals, oocyte release involves large-scale, cyclical tissue remodeling across five hormonally-controlled organs, which functionally degrade by mid-life. To better understand this system, we mapped the cellular compositional and transcriptional changes that occur during each estrus cycle phase, arising from earliest pregnancy, and upon aging at single-cell resolution in every organ of the mouse female reproductive tract. These data quantified the compositional transition between ovary, oviduct and uterus, which are dominated by stroma, and cervix and vagina, which are largely epithelial. Most importantly, our data provided unprecedented insight into how tissue remodeling by the stromal cells and inflammatory stimulus by the immune compartment can lead to the functional degradation of and susceptibility to disease within the female reproductive tract.

Our data showed that the adaptive immune cells are more prevalent in the upper tract compared to the lower reproductive tract which shows a profound shift towards innate immunity. Oviduct and uterus have an anti-inflammatory environment, dominated by wound-healing associated M2 macrophages (Madsen et al., 2013). In contrast, cervix and vagina have a pro-inflammatory environment, dominated by M1 macrophages consistent with potential microbial exposure (Zhou et al., 2018). We found that M1 macrophages were more numerous in the ovary, where they are required for folliculogenesis (Ono et al., 2018). We quantified how the immune compartment displays highly dynamic remodeling across the estrus cycle, with the more caudal organs such as vagina displaying stronger changes in the immune compartment. The lower FRT has recurrent acute inflammation during the cycle, contrasting with the persistent low-grade inflammation found in the ovary.

Previous experiments revealed that mouse uteruses infected during the estrus phase had significantly increased resistance to bacterial infection, compared to diestrus (Islam et al., 2016). Our data suggests a mechanism for this enhanced surveillance: we observed that during estrus the immune cell compartment expands in the uterus. More generally, the variation in immune cell composition and activation could explain why immune responses and protection of the reproductive system has been previously reported to vary along the cycle; indeed, vaccine-induced immunity is influenced by the cycle (Gallichan and Rosenthal, 1996).

Uniquely in the uterus, drastic ECM remodeling and cell proliferation recurrently occur in each cycle to prepare the endometrium for successful implantation and placentation (Kaloglu and Onarlioglu, 2010). In the absence of pregnancy, the proliferated tissue and secreted ECM are reabsorbed and degraded via scar-free remodeling mediated by fibroblasts (Bellofiore et al., 2018; Salamonsen et al., 2002). Our data allowed us to interrogate whether incomplete resolution of collagen deposition might contribute to development of organ fibrosis and thus reproductive senescence. Our single-cell transcriptomes across the cycle quantified ECM remodeling and stroma proliferation, which are particularly elevated in uterus and oviduct. By longitudinally profiling the accumulation of fibrosis across the FRT during aging, we discovered that the scale of ECM remodeling found in each organ during the cycle closely predicts fibrosis development in old age. Previous studies reported evidence of fibrosis development in post-menopausal endometrium (Jiménez-Ayala and Jiménez-Ayala, 2008; Noci et al., 1996); our data newly reveals that this fibrosis accumulates steadily across aging and that the age-related decrease in hormonal stimulation is not the primary driver of fibrosis development.

Within fibroblasts, aging also increased inflammation scores in all FRT organs, except ovary. The best predictor of an organ’s fibroblast inflammatory activity in old age was the amplitude of organ-wide inflammation during the cycle in young mice. This suggests a model whereby fibroblasts accumulate the memory of past inflammation, thus leading to age-related chronic inflammation.

In women, reproductive organs lose functionality faster than the somatic organs and display chronic inflammation and fibrosis by mid-life (Farage and Maibach, 2011; Noci et al., 1996). Our data suggest mechanisms that could explain the different organs’ susceptibility to cancer risk. Organs of lower FRT undergo recurrent cyclic acute inflammation and develop high-grade chronic inflammation in old age. Previous studies linked inflammation with cervical cancer progression (Mhatre et al., 2012). Our results revealed that the uterus was particularly susceptible to age-associated increases in inflammation and fibrosis. Endometrial cancer is the six most commonly occurring cancer in women worldwide (Ferlay et al., 2021). Elevated levels of pro-inflammatory cytokines pose a major risk in endometrial cancer development (Dossus et al., 2013), and a fibrotic microenvironment appears to contributes to endometrial tumor aggressiveness and drug resistance (Pradip et al., 2021). The risk of developing endometrial carcinomas may also be shaped by events that alter the number of menstrual cycles, such as oral contraception usage, number of pregnancies, and age of menarche and menopause (Havrilesky et al., 2013; Iversen et al., 2017; Michels et al., 2018). On balance, the fewer the number of cycles, the lower the cancer risk. This hypothesis could be directly addressed by reducing the number of estrus cycles, and evaluating the impact on FRT organ composition and inflammation. Age-related fibrosis and chronic inflammation development in the oviduct is comparable to the uterus. Inflammation and ECM-rich microenvironment were shown to directly contribute to seeding of cancer cells that originated in the oviduct to ovary thus causing high-grade serous ovarian cancer development (Alshehri et al., 2022; Jia et al., 2018).

Our work directly links intensity of inflammation and ECM activity during the estrus cycle and number of cycles with the severity of inflammation and fibrosis in old age. Our data supports a model wherein the incomplete resolution of inflammation and ECM remodeling during an increasing number of cycles adds to fibrosis inflammatory memory and ECM accumulation and lead to gradual development of fibrosis and chronic inflammation, predisposing organs to disease development.

## Acknowledgements and Funding

We thank the Institute of pathology (CMCP, Heidelberg University), the DKFZ Single-Cell Open Lab (scOpenLab), D. Krunic from the DKFZ microscopy core, F. Blum, K. Hexel, T. Rubner, S. Schmitt and M. Eich from the DKFZ flow cytometry and the animal caretakers from ATV.108 for technical support and assistance. We thank Roser Vento-Tormo, Camille Berthelot, Arnaud Krebs, Lenka Gahurová and Ludivine Doridot for offering suggestions and proof-reading the manuscript. This work was supported by NCT/Helmholtz core funding (B270 to D.T.O. and B210 to A.G.); European Research Council (788937 to D.T.O.); Helmholtz junior group leader post (to A.G.); DKFZ International Fellows Program (to A.T.); and the Deutsche Forschungsgemeinschaft (FI 2558/1-1 to A.G.).

## Author contributions

A.G., D.T.O. conceived the project and supervised the work.

I.W., A.T., A.G., D.T.O. designed experiments and wrote the manuscript.

A.T., N.S. M.L.K., J.P., K.D., F.G. performed the mouse handling, RNAscope, flow cytometry and sequencing experiments.

T.P. histopathological evaluation and grading.

I.W. performed computational analyses.

P.L. analyzed the Visium data.

K.D. performed the Visium experiments.

F.L. and A.G. performed supporting computational analyses.

F.L. curated data.

All authors had the opportunity to edit the manuscript, and all authors approved the final manuscript.

## Materials and Methods

### Mouse colony management

The C57BL/6 substrains J, N and Ly5.1 were obtained from Jackson Laboratories or Janvier. Females were maintained as virgins and housed in groups of up to six mice in Tecniplast GM500 IVC cages with a 12-hour light / 12-hour dark cycle. Mice had *ad libitum* access to water, food (Kliba 3437), and environmental enrichments. All colonies were regularly controlled for infections using sentinel mice to ensure a healthy status. All experiments were carried out in accordance with and approval of the German Cancer Research Center ethical committee and local governmental regulations (Regierungspräsidium Karlsruhe, animal license number DKFZ366).

### Estrus cycle staging cytology

Vaginal smears were collected using a pasteur pipette containing PBS and leaned towards or inserted in the vagina of the restrained mouse. Mucous tissue was then trickled on dry glass slides and stained by crystal violet staining solution (Sigma-Aldrich 61135) or panoptic staining (Carl Roth 6487.1). Cellular composition of the smears was analyzed according to known cell distribution patterns (Byers et al., 2012) using a ZEISS Discovery.V12 Stereoscope and images were acquired via a ZEISS Cell Observer® system with AxioCam MRc camera. If possible, smears were collected and analyzed from multiple consecutive days to better estimate estrus cycle course. On the day of tissue collection, estimation of the estrus cycle phase by smears was further complemented by the state of the vaginal opening and the thickness and vascularization of the uterine horns (Parker and Picut, 2016).

### Induction of decidualization

Three months old female C57BL/6J virgin mice were synchronized 3 days prior to mating by housing in cages containing bedding from C57BL/6 male mice. These females were allowed to mate with C57BL/6 males in one to one matings overnight. On the following morning, all plug-positive females were housed together and kept for 5 days (5.5 days post coitum) until sacrifice for organ harvesting. On average, two out of three plug-positive mice were pregnant at the day of sacrifice. Two uterine pieces, each enveloping an implanted embryo were removed per mouse and further processed for 10x or histology as described below.

### Histopathology and fibrosis quantification

After overnight fixation in 10% buffered formalin, representative specimens of the ovary, oviduct, uterus, cervix, vagina, and spleen were routinely dehydrated, embedded in paraffin, and cut into 4 μm-thick sections. All tissue sections were stained using a H&E standard protocol. In selected tissue sections, a Giemsa, Picrosirius Red, Congo Red, and AFOG (Acid Fuchsin Orange G) stain were performed according to respective standard protocols. To detect potential tumors, tumor classification immunohistochemistry was performed with anti-a smooth muscle actin (Abcam, ab5694), anti-desmin (ThermoFisher, RB-9014-Po), and anti-pancytokeratin (DAKO, Z0622). Whole-slide scans were acquired using the Aperio AT2 slide scanner (Leica) at 40x resolution. Raw image files are available from BioStudies (see data availability).

To quantify fibrosis, high-resolution whole tissue section images of Picrosirius red stained samples were acquired at 20x using a ZEISS Cell Observer® brightfield microscope and an AxioCam MRc camera. Fiji software (ImageJ ver. 1.53f51) was used to quantify percent of fibrotic area by setting a signal threshold in stroma-containing regions. RGB images were split into three channels. Signal quantification was performed on the green channel. Regions of interest were drawn around stroma areas and a threshold was set to correspond with the fibrotic area previously assessed by a pathologist. Collagen accumulations were defined as % of area with positive signal. Two replicates were used for 12, 15 and 21 month old mice, and three replicates were used for 3 and 18 month old mice. Analyses were performed independently by two authors (I.W. and A.T.) to reduce stroma area selection bias. The macro is available from the GitHub repository (Code availability). Technical replicates or quantified regions of interest from the same sample were averaged and treated as one biological replicate in a linear regression model.

### RNA in situ hybridization

To detect and quantify Collagen, Type 1, alpha 1 (Col1a1, ACD 319371) and Epithelial Cell Adhesion Molecule (Epcam, ACD 418151-C2) mRNA, an ISH was performed using the RNAScope Multiplex Fluorescent V2 Assay (ACD 323100) with Opal fluorophore reagents (Akoya Biosciences) according to the manufacturers’ instructions. In brief, fresh and fixed frozen samples were collected according to manufacturers’ protocol and cut at 10µm thickness using a Leica CM 3050S cryotome. Target probes (Col1a1, 319371-C1; Epcam, 418151-C2) were applied to the sample and baked at 40°C for 2h. Opal dyes 520 (FP1487001KT),570 (FP1488001KT) and 690 (FP1497001KT) were applied at a 1:1000 to 1:750 dilution and counterstained with DAPI. Images were taken using the ZEISS Cell Observer® fluorescence microscope and a ZEISS AxioCam MRm camera at 20x resolution.

Collagen signal was quantified as percent area of total DAPI area using ImageJ. Samples which were run with negative control probes (ACD 320871) were used to subtract background signals beforehand. The macro for assessing signal thresholds is available from GitHub (Code availability).

### Tissue collection and preparation

All four phases of estrus cycle in ovary, oviduct, uterus, cervix, vagina, and spleen were collected in triplicate from 3 and 18 month old mice. Seven replicates were collected for decidualized tissues. Additional biological replicates for samples that failed QC requirements were generated as deemed necessary.

Samples “Ind001-vagina06” and “Ind001-uterus07”, “18mo_Ind001-ovary01”, “18mo_Ind001-spleen01” and “18mo_Ind001-ovary02” were excluded from the analysis due to problematic sample preparation (Table S1). Instead, additional mice were sacrificed to collect the vagina in proestrus (“Ind001-vagina13”), uterus in diestrus (“Ind001-uterus16”), ovary from 18 months old mice (“18mo_Ind001-ovary04”, “18mo_Ind001-ovary05”) and spleen “18mo_Ind001-spleen05”.

Reproductive tract organs and spleen were collected from mice immediately following cervical dislocation. All organs were manually dissected using a ZEISS Discovery.V12 Stereoscope to remove surrounding fat and connective tissue. Samples were then either processed by enzymatic digestion for single cell sequencing, fixed in 10% formalin for FFPE-histology, or fixed and slowly frozen in O.C.T. Medium (ThermoFisher) for cryo-histology.

### Generation of single cell suspensions

To generate single cell suspensions, freshly isolated whole organs including ovary, oviduct and tissue pieces from uterus, cervix, vagina and spleen were treated by enzymatic digestion. All tissues were initially incubated separately in 2 ml Eppendorf tubes containing 600 µl of 0,25% trypsin in HBSS and digested at 37°C with gentle rocking. After 30 minutes, 600µL of a second digestion buffer containing Collagenase I (1.25 mg/mL), II (0.5 mg/mL), IV (0.5 mg/mL), and Hyaluronic acid (0.1 mg/uL) in HBSS was added for another 2 hours. After quenching the digestion by adding 600µL HBSS with 10% FBS, the cell suspensions were passed through a 40µm cell strainer (Greiner Bio One) to remove cell debris and buffer residue. Cell suspensions were gently centrifuged at 350g for 8 min at 4°C. Cells were resuspended in PBS containing 0,04% BSA, 1 mM EDTA and propidium iodide (PI) was added to final concentration of 1 µg/ml prior to fluorescence-activated cell sorting (FACS). Larger cells such as oocytes and smooth muscle cells were excluded in the cell straining step.

### Flow cytometry staining and acquisition

After dissection and digestion, cells were filtered through 40 µm cell strainers (Falcon), followed by washing and centrifugation for 5 min at 250 g at 4°C. For flow cytometric analysis, cells were resuspended in 20 µl PBS plus Viability Dye eF506 (eBioscience, conc. 1:500) and incubated for 10 min at RT in the dark. Proceeding with cell staining, 100 µl PBS plus 2% FCS with corresponding antibodies (see Table 1) was added and cells were incubated for 30 min at 4°C in the dark. Post staining, cells were washed again and analyzed using the BD LSR II cytometer according to manufacturer’s instructions and marker combinations and gating strategies (Figure S3b).

**Table 1:** Anti-mouse antibodies / Hematopoietic cells panel.

| Antigen:Fluorophore | Antibody dilution | Clone | Company |
| --- | --- | --- | --- |
| F4/80:PB | BV421 (F4/80) – 1:100 | MF48028 | Invitrogen |
| NK1.1:BV785 | BV785 (NK1.1) – 1:700 | PK136 | BioLegend |
| CD11c:PE-Cy7 | PECY7 (Cd11c) – 1:300 | N418 | BioLegend |
| CD11b:APC | APC (Cd11b) – 1:1000 | M1/70 | eBioscience |
| CD45:AF700 | AF700 (Cd45) – 1:200 | 30-F11 | Invitrogen |
| Gr1:APC-eF786 | APCCy7 (Gr1) – 1:500 | RB6-8C5 | eBioscience |

FAC sorting as preparation for single-cell transcriptional analysis was performed on the FACSAria II from BD Biosciences using an 85µm nozzle. Gating of live cells was achieved by exclusion of PI-high cells. Doublets were excluded by plotting SSC width versus SSC area. Approximately 70.000 cells were collected in sorting media (containing 0.04% BSA and 1mM EDTA in PBS) in 1.5 ml Eppendorf tubes, chilled on ice, and immediately processed for single-cell transcriptional analysis.

### Generation of single cell transcriptomes

Mouse reproductive tissues and spleen were enzymatically dissociated and FAC sorted, and the undiluted single-cell suspension at a concentration of 467 cells/µl was loaded per channel of either the ChromiumTM Single Cell B or G Chip (10X Genomics® Chromium Single Cell 3’ Reagent Kits v3.0 and Chromium Next GEM Single Cell 3’ Reagents Kits v3.1, respectively), aiming for a recovery of 5,000 cells. Reverse transcription and library construction were carried out according to the manufacturer’s recommendations. Libraries were sequenced on Illumina NovaSeq 6000 using a paired-end run sequencing 26 bp on read 1 and 98 bp on read 2 (Table S1).

### Spatial transcriptomics

12-14 weeks old *Mus musculus* (C57Bl6/J) female mice were sacrificed at diestrus. Ovaries were dissected and embedded into an optimum cutting temperature matrix (O.C.T., Tissue Tek) using a bath of pre-cooled (−60-70°C) isopentane (Sigma) on dry ice. Blocks were cut using cryo-microtome (CM3050 S, Leica), head temperature set at -10°C. 10 µm thick tissue slices were placed on Visium Spatial Gene Expression Slides (10X Genomics) and stained with Hematoxylin and Eosin (H&E) with reduced time to 5 min for hematoxylin and 30s for blueing agent. Libraries were prepared by manufacturer’s recommendations, using Dual Index Kit TT Set A (10X Genomics) for indexing. Samples were sequenced on NovaSeq6000.

### Computational quality control, normalization, cell type annotation and batch correction

Raw sequencing reads were processed using Cell Ranger analysis pipeline (v 3.0.1). The “cellranger count” command was used to generate filtered and raw matrices. Reads were aligned against the mouse genome version mm10 (Ensembl release 93). Filtered gene-barcode count matrices were further analyzed using the R package Seurat (Hao et al., 2021).

To remove low quality cells, an adaptive filtering threshold approach was used based on high mitochondrial RNA content, extreme numbers of counts (count depth), and extreme numbers of genes per barcode. Cells were filtered based on the median absolute deviation (MAD) from the median value of each metric across all cells. Specifically, we considered a value as an outlier when differing by more than 3 MADs from the median. The filtering step was performed using the R package Scater (McCarthy et al., 2017). Counts were normalized using the ScTransform normalization approach of Seurat. Cell cycle gene effect was regressed out using the CellCycleScoring function in Seurat. All clusters in all samples showed consistently low doublet scores using doubletCluster and doubletCells of R package Scran (Lun, 2016).

Each organ was processed independently for cell type annotation. Organ-specific UMAPs were constructed using a subset of features (genes) exhibiting high cell-to-cell variation which were identified by modeling the mean-variance relationship. The top 2000 features were used to perform PCA analysis. To cluster the cells, a K-nearest neighbor (kNN) graph based on the euclidean distance in PCA space was first constructed using the first 30 PC components as input. Next, the Louvain algorithm was applied to iteratively group cells. We identified the cell types in each cluster using a combination of manual and automated approaches from known marker genes (Table S2). First, clusters were assigned to known cell populations using cell type–specific markers obtained through the FindAllMarkers function. Multiple testing correction was performed using Benjamini-Hochberg procedure. Second, the R package Garnett (Pliner et al., 2019) in cluster extension mode was used to annotate cells in a semi-automated manner. Because some clusters remained unclassified by either the manual or semi-automated approach - or in rare cases were differently classified by the two approaches - a Support vector Machine with rejection (SVMrej) was applied as an additional automated classifier. Cluster annotations in agreement between the manual and automated approach were used as the training set for the SVMrej. The e1071 library was used to implement the SVMrej classifier. Classification was performed using a linear kernel with the cost function set to 10. Rejection rates of 10% and 30% were used to classify level 1 and level 2 annotations, respectively. Cell clusters of 18 month old and pregnant mice were annotated using an SVMrej classifier trained on cell clusters of normally cycling young mice.

Cells from multiple organs in the estrus cycle were integrated together for visualization purposes. Integration and batch correction of samples of young cycling mice was performed using the Reciprocal PCA together with “LogNormalize” normalization method from Seurat. For each organ, in order to remove batch effects we chose the sample with the highest number of cells (regardless of its cycle phase) to anchor pairwise comparisons. Integration of young (diestrus) and old mice followed a similar approach.

Integration of samples of pregnant mice with metestrus samples was performed separately using Canonical correlation analysis together with SCT normalization (Seurat). Batch corrected data was used only for UMAP visualization purposes; all other downstream analyses including differential expression were performed on uncorrected data. To improve the visualization of gene expression in UMAP plots, we used a kNN-pooling approach (Frauhammer, Felix and Anders, Simon, 2022). A kNN graph was constructed based on euclidean distance in PCA space, and cell counts were pooled across 50 nearest neighbor cells, thus decreasing technical noise in scRNA-seq caused by dropout events.

### Differential cell abundance analysis

To assess if the proportions of cell populations in individual organs change along the estrus cycle and in aging, a compositional regression model was used. Cell population compositions were used as dependent variables, and estrus cycle phases or age as independent variables. Estrus cycle phases were compared in a pair-wise manner. Components of the compositions were amalgamated and expressed as compositions of two components due to low number of replicates. Prior to model fitting, composition components were transformed using isometric Log-Ratio Transformation. The R package Compositions was used to perform the Log-Ratio transformation and compositional regression. Innate vs adaptive immune cells ratio was calculated as ratio of number of innate immune cells (N, DC, M1Mp, M2Mp, Mp and MaC) and number of adaptive immune cells (NKC, BC, iNkT, MAIT, MTC) in each individual.

### Optimal transport

Balanced optimal transport (OT) analysis was performed to assess the magnitude of transcriptional changes between cell populations of all organs in different phases of the estrus cycle, as well as old and young (diestrus) cell populations. For specific subgroups of samples, we performed simultaneous NMF embeddings; these two subgroups included all organs of all estrus phases and all organs of young (diestrus) together with all organs of old mice. NMF embeddings were calculated on the set of top 2000 most highly-variable genes using the Block Principal Pivoting method of the PLANC library (Ramakrishnan et al., 2016). Rank 10 of NMF embeddings, chosen based on decrease in cophenetic coefficient (Brunet et al., 2004), was used for OT distance calculation. OT distances were calculated for all cell populations which contained at least 100 cells in any of the compared groups. As the balanced optimal transport problem is constrained with a mass balance condition, the OT distance between two cell populations was calculated as an average of 100 random samples of 100 cells in a stochastic sampling approach. The OT distance was defined as a minimum-cost flow solution problem and was solved using Munkres algorithm (Kimmel et al., 2019).

### Differential gene expression (DGE) analysis

DGE analysis was performed using a multi-level generalized negative binomial regression model with a random intercept. Normalized gene counts were used as the dependent variable, while estrus cycle phases or age were used as the independent variable, and sample label as random effect. The model was fitted gene-wise for each cell subpopulation. The cycle phases were compared in a pair-wise manner. The model fitting was performed using the glm.nb function of the lme4 R package. Only genes that were expressed across 10 cells with at least 1 count were used in model fitting. If the model fitting showed a singular fit (indicating overfitting) the p-value was set to *NA*. P-values were corrected for multiple testing using Benjamini-Hochberg procedure. All genes with corrected p-values smaller than 0.05 were considered differentially expressed.

### Overrepresentation analysis (ORA)

A hypergeometric test was used to perform ORA analysis. Gene sets used in ORA analysis are part of the MsigDB (**https://www.gsea-msigdb.org/gsea/msigdb/**) pathway collection (H; C2: Kegg, Reactome; and C5: GO Biological Process). For the ORA in figure 3, we used MSigDB H and Reactome, and excluded pathways with the term “Disease” in their descriptor. Multiple testing correction was performed using the Benjamini-Hochberg procedure.

### Scoring of gene set activity in single-cell RNA-seq data

Scoring of gene set activity was performed using the AUCell R package (Aibar et al., 2017). AUCell was used to assess if certain gene sets were enriched within the top 5% expressed genes for each cell. Gene sets used in the analysis are the same as gene sets used in ORA. For the scoring of the activity of target genes of each ligand in fibroblasts, we used the NicheNet ligand-target model to obtain a list of predicted targets for each ligand (Browaeys et al., 2020). AUCell scores were calculated for each cell and averaged across conditions for each cell population. Average scores across the cycle were weighted to account for the different phase lengths (the cycle was roughly estimated to be partitioned 60% diestrus and the remainder equally divided between proestrus, estrus and metestrus, (Byers et al., 2012).

### Genes expression association with cycle phase using Mutual-information (MI) and comparison to human endometrium dataset

Human endometrium dataset was retrieved from NCBI’s Gene Expression Omnibus (accession code **<u>GSE111976</u>**). Raw count matrices of Fluidigm C1 dataset were normalized using the ScTransform normalization approach. Cycle phase labels of human samples were assigned based on the original publication’s metadata (Wang et al., 2020).

Dependence of gene expression and cycle phase label in fibroblasts and decidual cells was calculated as the MI between these two variables, as described in (Wang et al., 2020). The same calculation was performed for the human fibroblasts. Briefly, for each gene, MI was calculated using the Java implementation of ARACNe-AP (Lachmann et al., 2016). The statistical significance of MI was evaluated using the permutation approach, in which MI value for each gene was compared to a null distribution obtained by permuting cycle phase labels 1,000 times. Multiple testing correction was performed using Benjamini-Hochberg. As the mouse dataset was substantially bigger than the human dataset, MI and its associated p-value for the mouse dataset were calculated on 100 random samples of 2000 cells in a stochastic sampling approach. P-values associated with genes in all mouse random samples were aggregated using Fisher’s method (R package aggregate). Genes associated with p-value smaller than 0.05 were considered as cycle-associated genes in mouse and human.

We identified the set of conserved transcriptional changes between human and mouse cycles by identifying homologous cycle-associated genes that showed the same directionality of regulation in comparison to adjacent cycle phases in paired mouse-human cycle phases. For instance, a mouse gene upregulated in proestrus compared to diestrus and the homologous human gene upregulated in proliferative early compared to secretory late. Genes that showed opposite directionality of regulation (e.g. up-regulation in humans, down-regulation in mice) were considered divergent. Mouse and human cycle phases were paired based on ovulation timing and uterine cycle events (proliferation vs secretion) (Ajayi and Akhigbe, 2020; Greaves, 2012). Proestrus was paired with proliferative early phase, estrus with proliferative late, metestrus with secretory early, decidualization with secretory mid and diestrus with secretory late. Cells of menstruation phase in humans could not be paired with normally cycling mouse cells and were excluded from this analysis. To calculate the proportion of the conserved genes expected by chance in each cycle phase, labels of upregulated, downregulated and neutral (not up- or down-regulated) genes of all homologs in mouse and human for corresponding cycle phases were permuted 100 times and proportions of the conserved/divergent genes were calculated per cycle phase. Final value of conserved genes proportion was calculated as an average of proportion of conserved downregulated and upregulated genes in all permutations runs.

### Cell-to-cell communication analysis

To assess cellular communication from different cell types to fibroblasts, we used a multiplication of expression between receptors and ligands (expression product) as a communication score. The list of mouse ligand-receptor pairs that was used in the analysis was extracted from CellChat (Jin et al., 2021) and CelltalkDB repositories (Shao et al., 2021). To compare cellular crosstalk among the different organs, we first focused on the average expression values of ligands in all cell types and receptors in fibroblasts, regardless of the source of the ligand. To calculate the communication score, averaged ligand expression counts in all cells from all cycle phases were multiplied with averaged receptor counts in fibroblasts. For multi-subunit receptors, the subunit with the minimum average expression was used in our calculations as previously proposed (Efremova et al., 2020).

The statistical significance of the difference of expression product in all reproductive organs was evaluated using a permutation approach. All pairwise combinations of log ratios of expression product in all organs for each ligand-receptor pair were compared to null distribution obtained by permuting the organ labels 1000 times. A similar approach was used to evaluate the statistical significance of the difference of expression product upon aging. For each organ, log ratios of expression product of old and young fibroblasts in diestrus were compared to null distribution obtained by permuting the age labels 1000 times. Multiple testing correction was performed using Benjamini-Hochberg procedure.

For selected ligand-receptor pairs, we then determined which cell type was the likely source of the ligand. The number of ligand counts that each cell of respective cell type produced was calculated, thus taking into consideration the cell abundance and average expression of ligand in each cell type. We chose to closely analyze the ligand-receptor pairs related to inflammation and ECM, based on the known role of receptor-ligand interactions in shaping their functions (Gurtner et al., 2008; Koliaraki et al., 2020; Turner et al., 2014).

### Spatial cell-to-cell communication analysis

To validate the cell-to-cell communication results, we generated spatial transcriptomics data using Visium (10X Genomics). The raw reads were processed using spaceranger (10X Genomics, v1.3.1). The data was then analyzed using Seurat (v4.0.3 in R 4.0.0) (Hao et al., 2021) and raw counts were corrected using the SCT method.

For each spot, the neighborhood refers to the combination of the spot itself and the directly adjacent spots. All spots containing stromal cells (corrected counts for Col1a1 above 1) and having stromal cells, macrophages (expression of Cd68), or granulosa cells (corrected counts for Serpine2 above 3) in their neighborhood were considered. Communication between spots containing stromal cells and ligand source cells was considered possible if: 1) the stromal spot expressed any of Tgfbr3 or Tgfbr1+Tgfbr2, and 2) Tgfb1 and the correct cell type marker were expressed in any of the neighborhood spots.

### Single-Cell Regulatory Network Inference and Clustering (SCENIC) analysis

pySCENIC was used to perform single-cell regulatory network analysis in fibroblasts (Van de Sande et al., 2020) by using normalized gene expression values of specific subsets of cells, i.e. fibroblasts of all organs in all phases of the cycle, fibroblast of all organs in diestrus and aged samples, and uterine fibroblasts in metestrus and pregnant samples. The gene co-expression networks were determined using grnboost2, enriched transcription factor motifs were predicted using ctx function and regulon activity scores were calculated using AUCell. To assess if regulons are differentially active across FRT organs or in aging, the activity score for each regulon was compared to null distributions obtained by permuting organ- or age-labels 1000 times. Multiple testing correction was performed using Benjamini-Hochberg procedure.

Selection of the subset of transcription factors related to inflammation and ECM was based on the overlap and enrichment of transcription factor target genes and target genes of selected ligands obtained from the NicheNet database. Additionally, classification of transcription factors as fibrosis and/or inflammation associated was based on the enrichment score of transcription factor target genes in inflammation (Hallmark collection) and ECM organization (Reactome collection) pathways. Transcription factor target genes were identified in pySCENIC analysis. Jaccard index was used to quantify overlap between target genes and pathway related genes, and a hypergeometric test was used to assess the enrichment.

### Single-cell trajectory inference

Slingshot was used to infer fibroblast trajectories along the estrus cycle (Street et al., 2018). Linear discriminant analysis (LDA) was used to perform dimensionality reduction. LDA was performed on cycle-associated genes determined in MI approach using the mda R package. We clustered the cells using k-Means, and we fitted the principal curve through fibroblast clusters using the Slingshot function.

### Differential cell heterogeneity analysis

Differential Shannon Entropy (ShE) was used to assess the differences in transcriptional heterogeneity between young (diestrus) and old cell populations. Differential ShE was calculated using the EntropyExplorer package (Wang et al., 2015). Multiple testing correction was performed using Benjamini-Hochberg procedure.

### Linear mixed models of inflammation

To estimate the rate of inflammaging in the different organs, we fitted a linear mixed model at the cell level including age, organ and the interaction of age and organ as fixed effects, and individual (mouse) as a varying intercept random effect.

### Data availability

All sequencing data and expression count matrices are deposited in Arrayexpress under accession number E-MTAB-11491. Imaging raw and processed data, as well as flow cytometry raw data together with the MIFlowCyt protocol are available at Biostudies (S-BIAD482, S-BIAD476 and S-BSST864). Dynamics of cellular abundance, gene expression, cell-to-cell communication and transcription factor regulatory networks in FRT organs in estrus cycle, decidualization and aging, can be explored through an interactive tool at **https://cancerevolution.dkfz.de/estrus/**

### Code availability

Code used in this study is available in github repository: https://github.com/goncalves-lab/estrus_cycle_study.

