## Supplementary Figures for "The function and decline of the female reproductive tract at single-cell resolution"

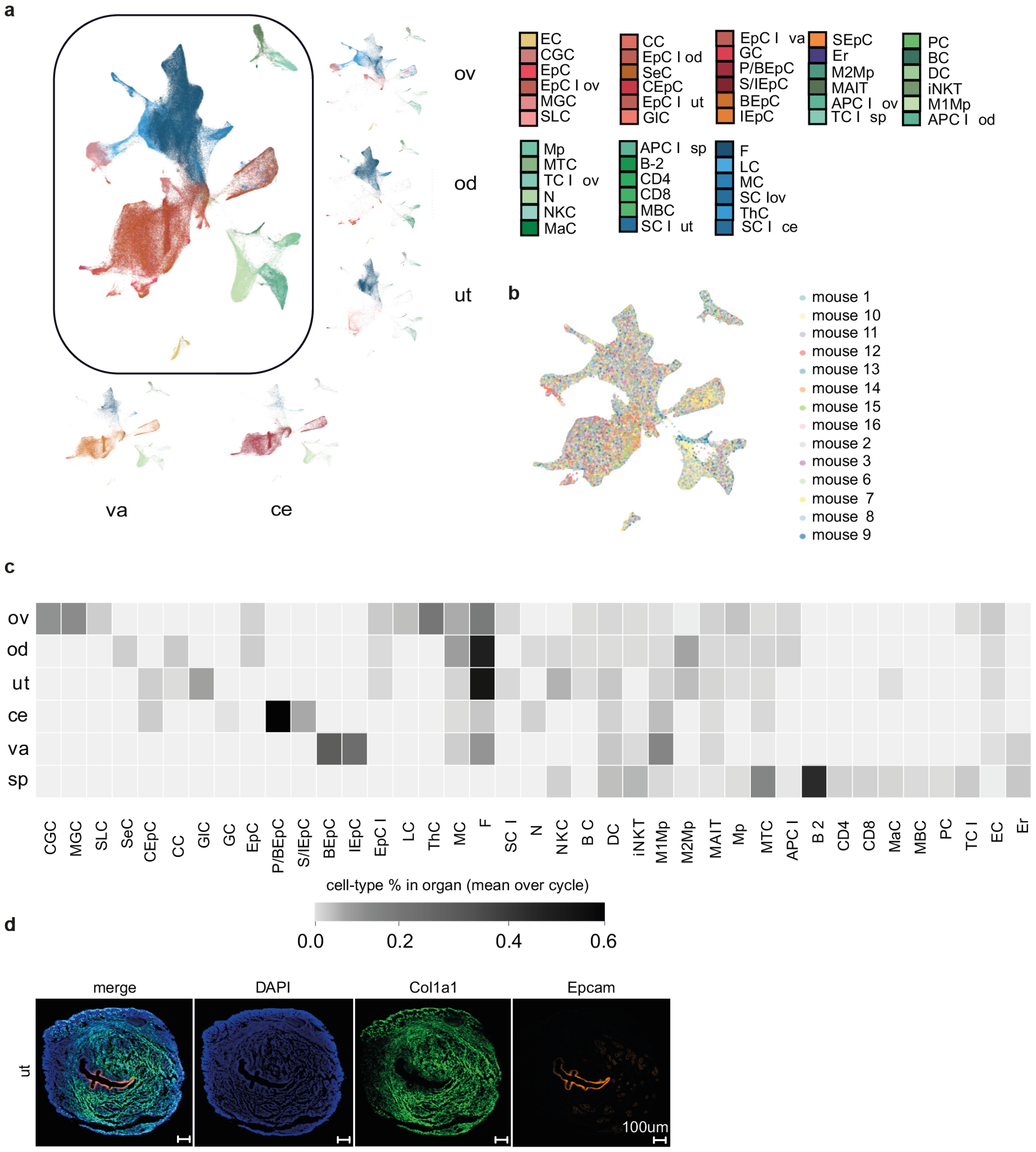


**Figure S1. (a)** UMAP plot of the young cycling mouse cells. Full names of assigned cell type abbreviations are listed in Figure 1c. **(b)** UMAP plot of young cycling mouse cells. Cells are colored based on individual mouse. Uniform mixing of cells originating from different mouse individuals indicates absence of batch effect. **(c)** Proportional heatmap of the most abundant cell types by organ in young cycling mice (a subset of this heatmap is shown in Figure 1c). **(d)** Expression of Col1a1 and Epcam in young uterus (diestrus) detected using RNA in situ hybridization.

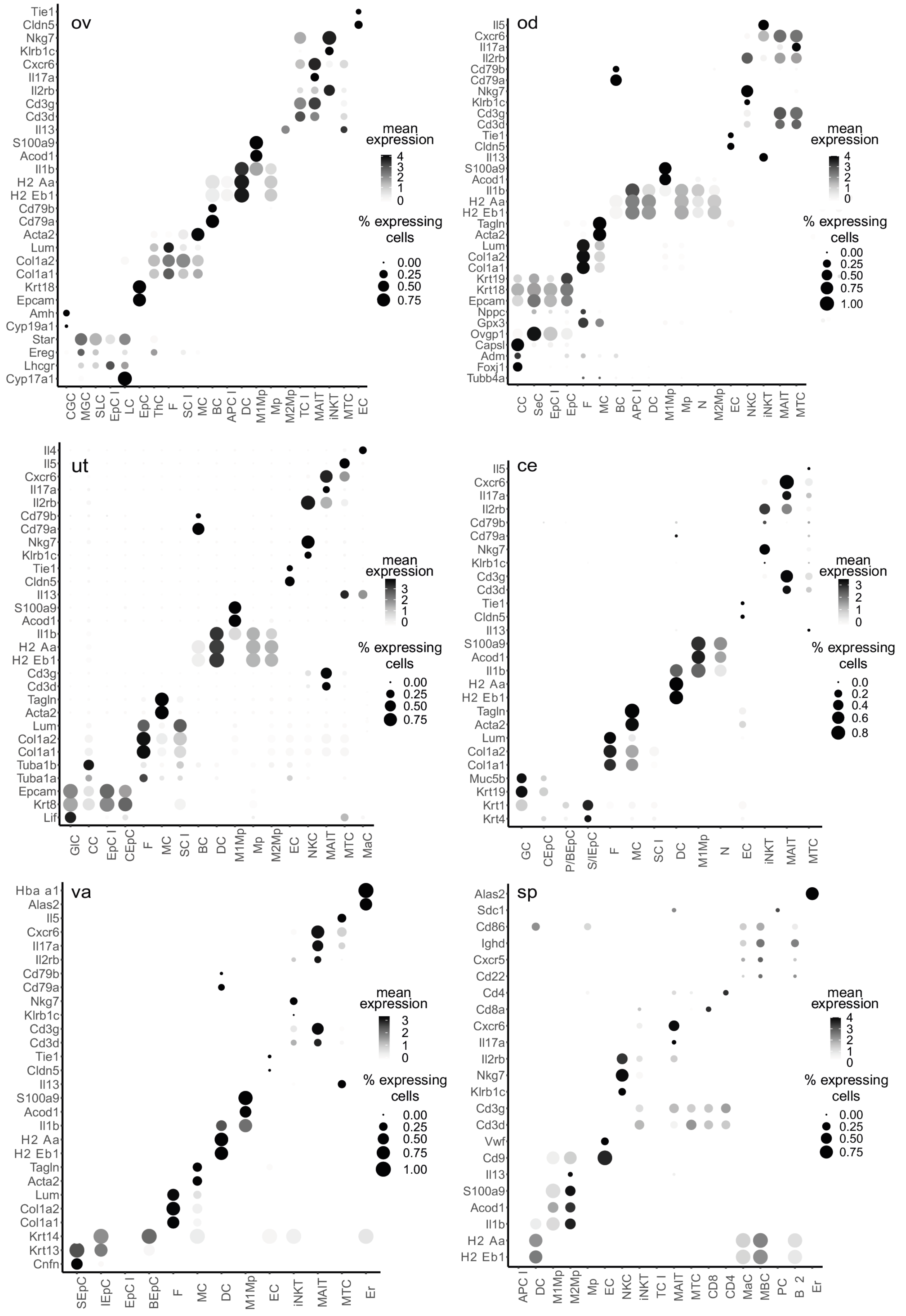


**Figure S2.** Marker genes used in the classification of the ovary, oviduct, uterus, cervix, vagina and spleen cell types.

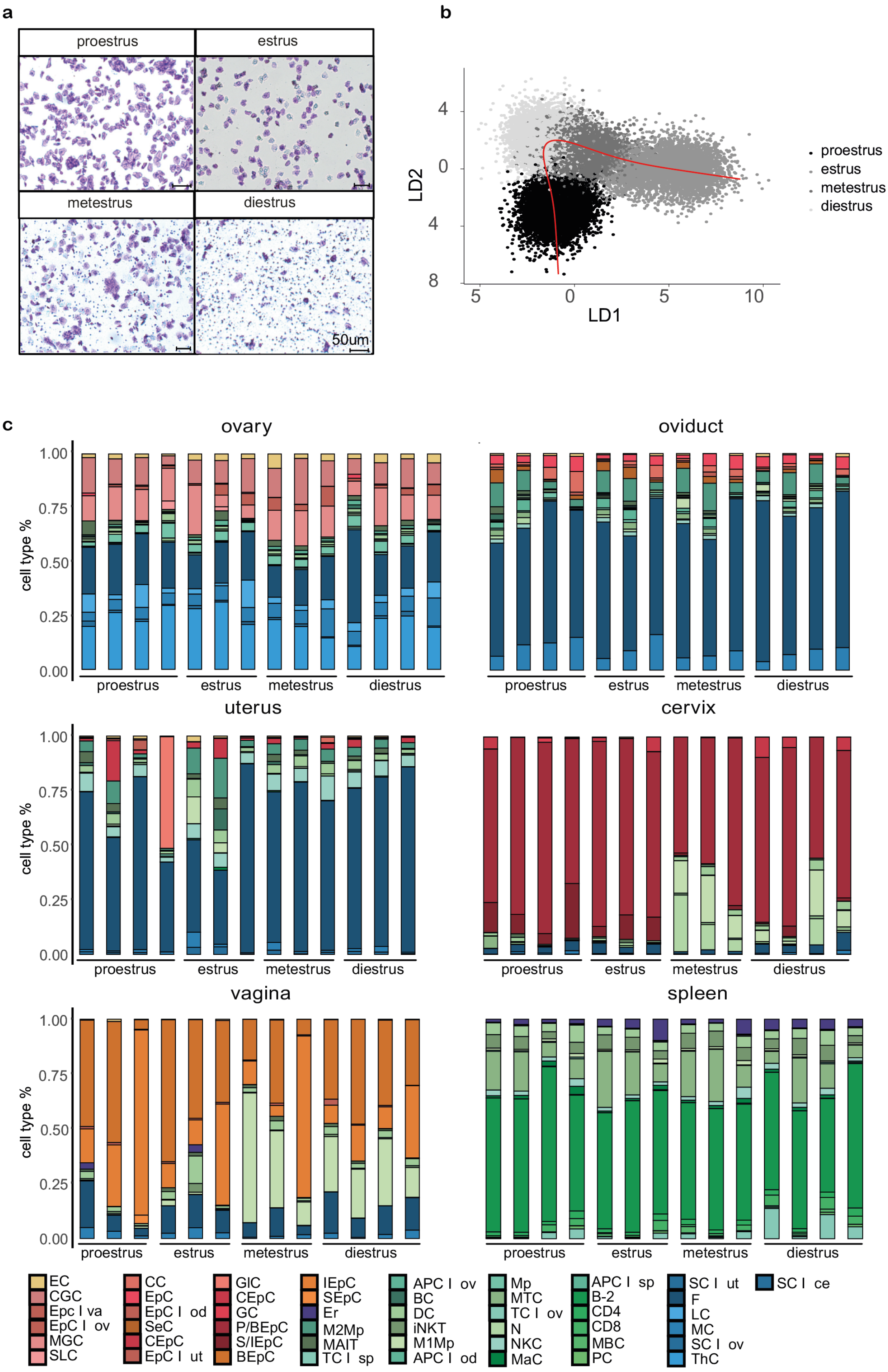


**Figure S3. (a)** Crystal violet staining of vaginal smears of young cycling mice. Cycle phases (proestrus, estrus, metestrus and diestrus) were assigned based on occurrence of leukocytes, and nucleated and cornified epithelial cells. **(b)** Pseudotime-time trajectory of uterine fibroblasts across the cycle. Cell clusters are colored according to cycle phases. Dimensionality reduction was performed using LDA and cell features projected to the first two LD components were plotted. **(c)** Barplots showing % of each cell type in ovary, oviduct, uterus, cervix, vagina and spleen at each phase of the cycle in each biological replicate.

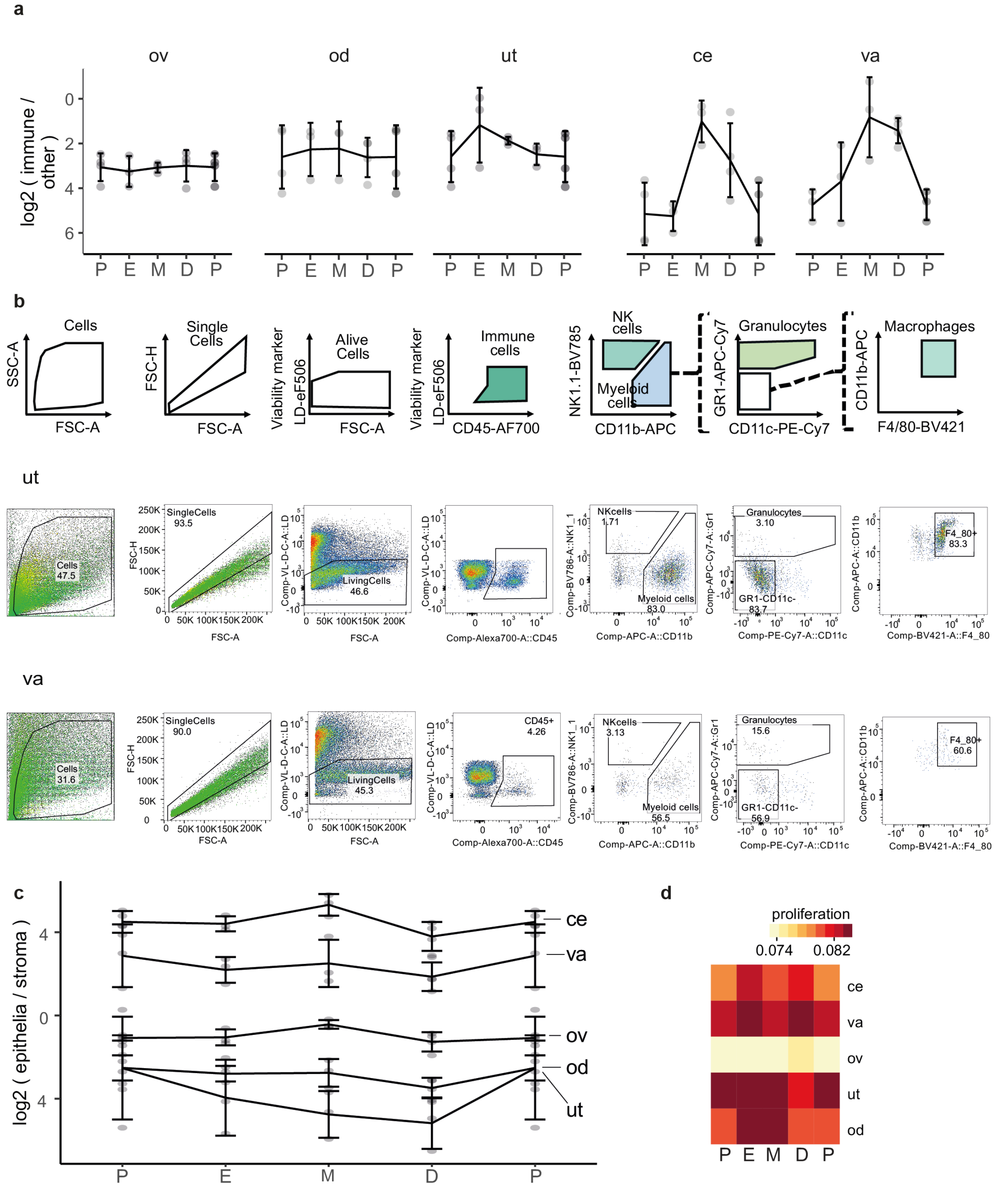


**Figure S4. (a)** The ratio of immune to other cells for each biological replicate (dot) is plotted. The average ratios (line) are shown together with their standard errors (vertical error bars). **(b)** FACS gating strategy used to quantify immune and myeloid cells in uterine and vaginal samples across the cycle. **(c)** The ratio of epithelia to stroma for each biological replicate is plotted. **(d)** Average activity score of genes promoting cell proliferation (GO:0008284) calculated in stromal cells using AUCell.

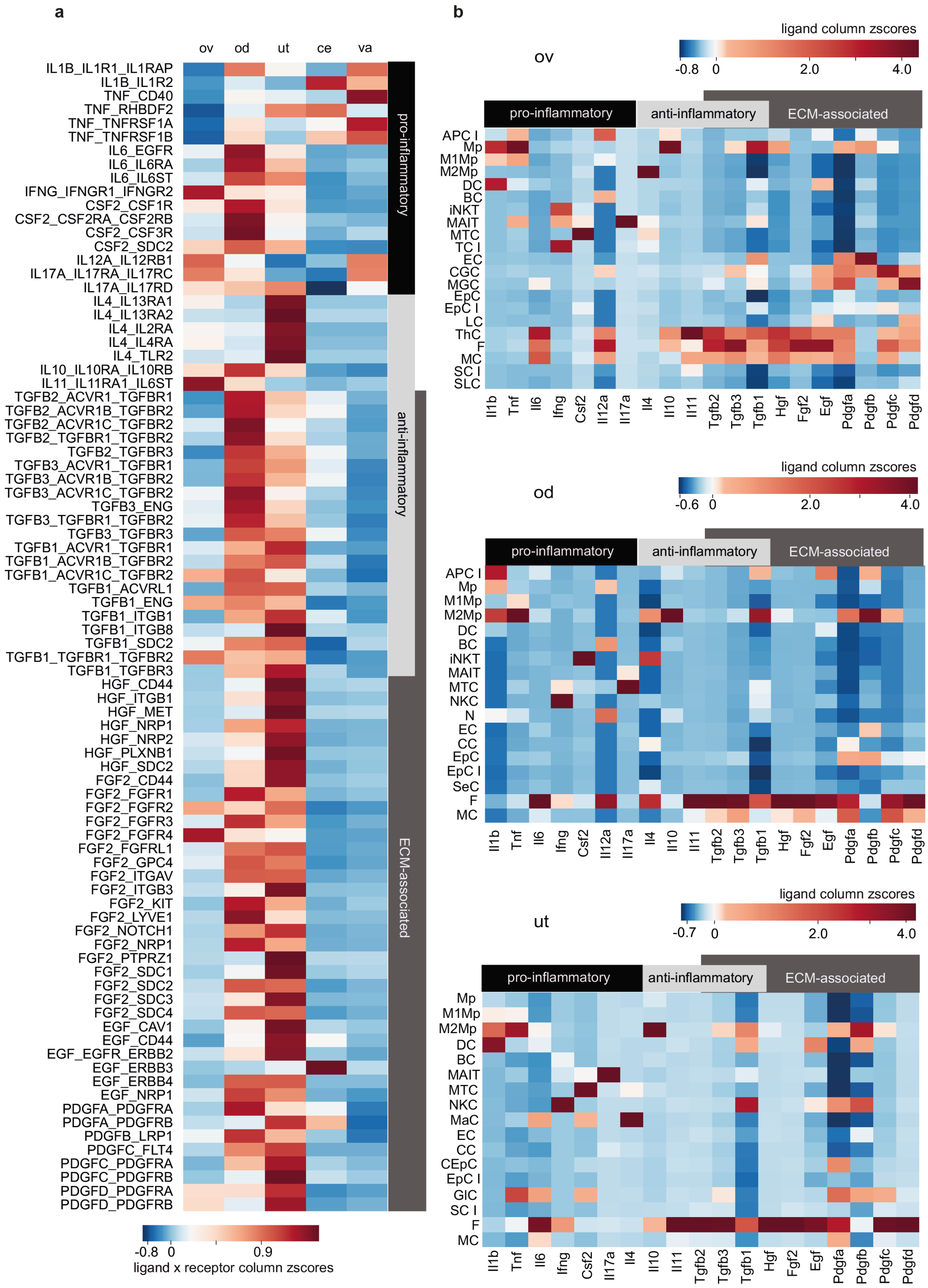


**Figure S5. (a)** Heatmap showing the z-scores of ligand-receptor products averaged across phases. Subset of this heatmap is shown in Figure 3e. Ligand expression is averaged across all cell types; receptors are in fibroblasts only. **(b)** Heatmap showing ligand expressions in all cell types of ovary, oviduct and uterus.

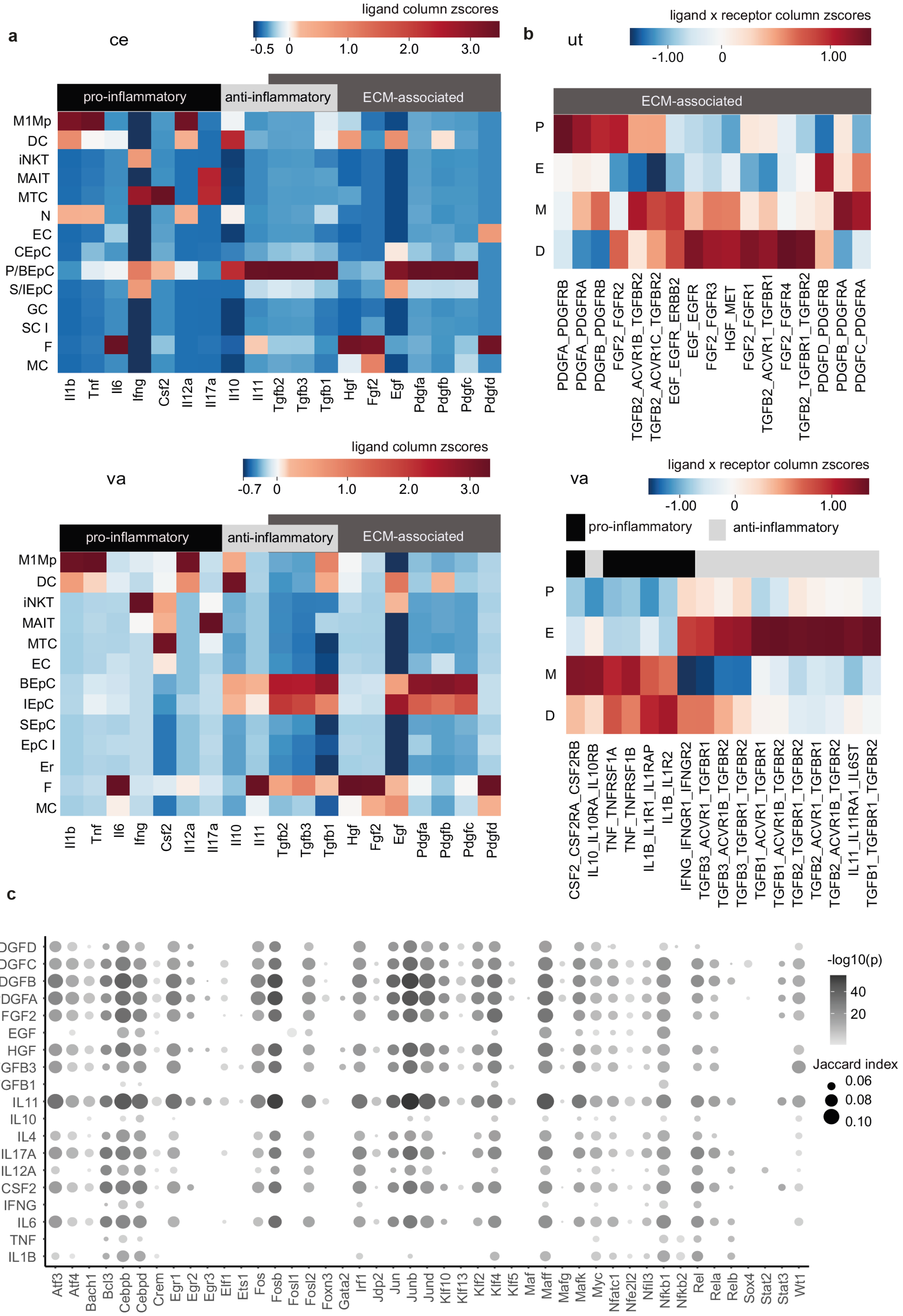


**Figure S6. (a)** Heatmap showing ligand expressions in all cell types of cervix and vagina. **(b)** Heatmap showing the z-scores of ECM-associated ligand-receptor products in uterus and inflammatory ligand-receptor products in vagina across cycle phases. Ligand expression is averaged across all cell types; receptors are in fibroblasts only. **(c)** Enrichment and overlap of target genes of ligands and target genes of transcription factors active in fibroblasts. Enrichment was determined using hypergeometric test and overlap using Jaccard index.

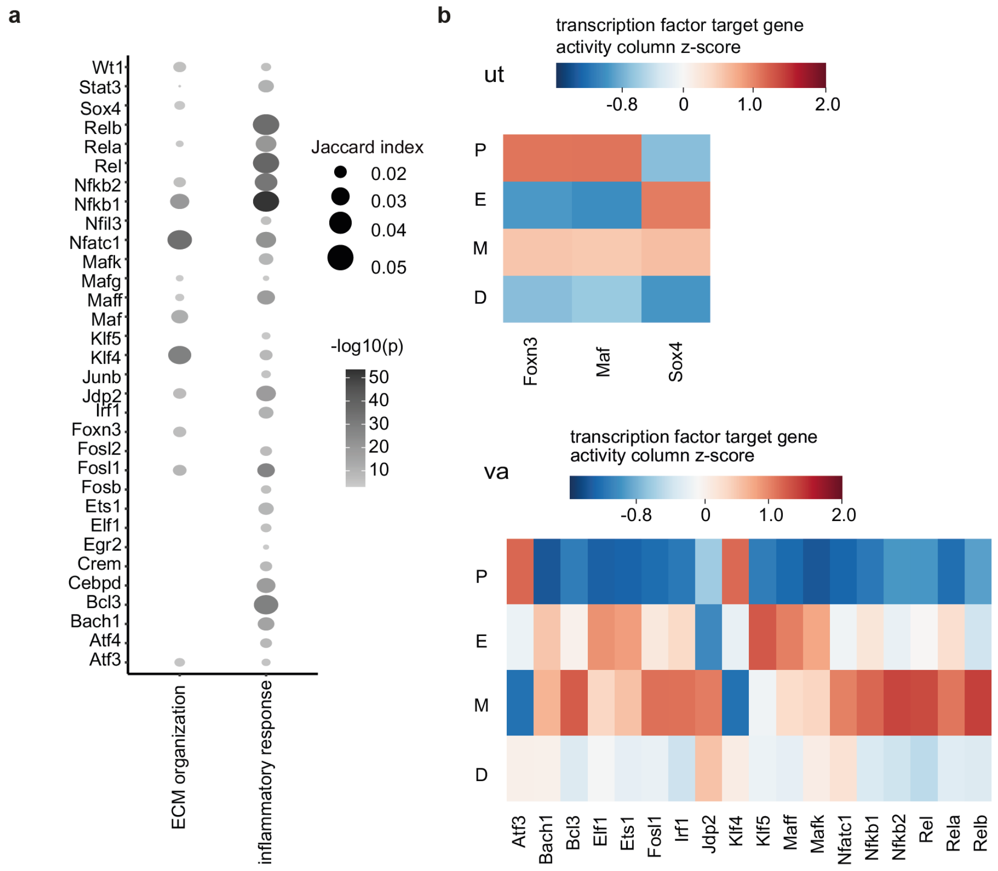


**Figure S7. (a)** Enrichment and overlap of target genes of transcription factors active in fibroblasts and genes in ECM organization and inflammation pathways. Enrichment was determined using hypergeometric test and overlap using Jaccard index. **(b)** Activity scores of targets of transcription factors associated with ECM regulation in uterine fibroblasts and inflammation in vaginal fibroblasts across the cycle.

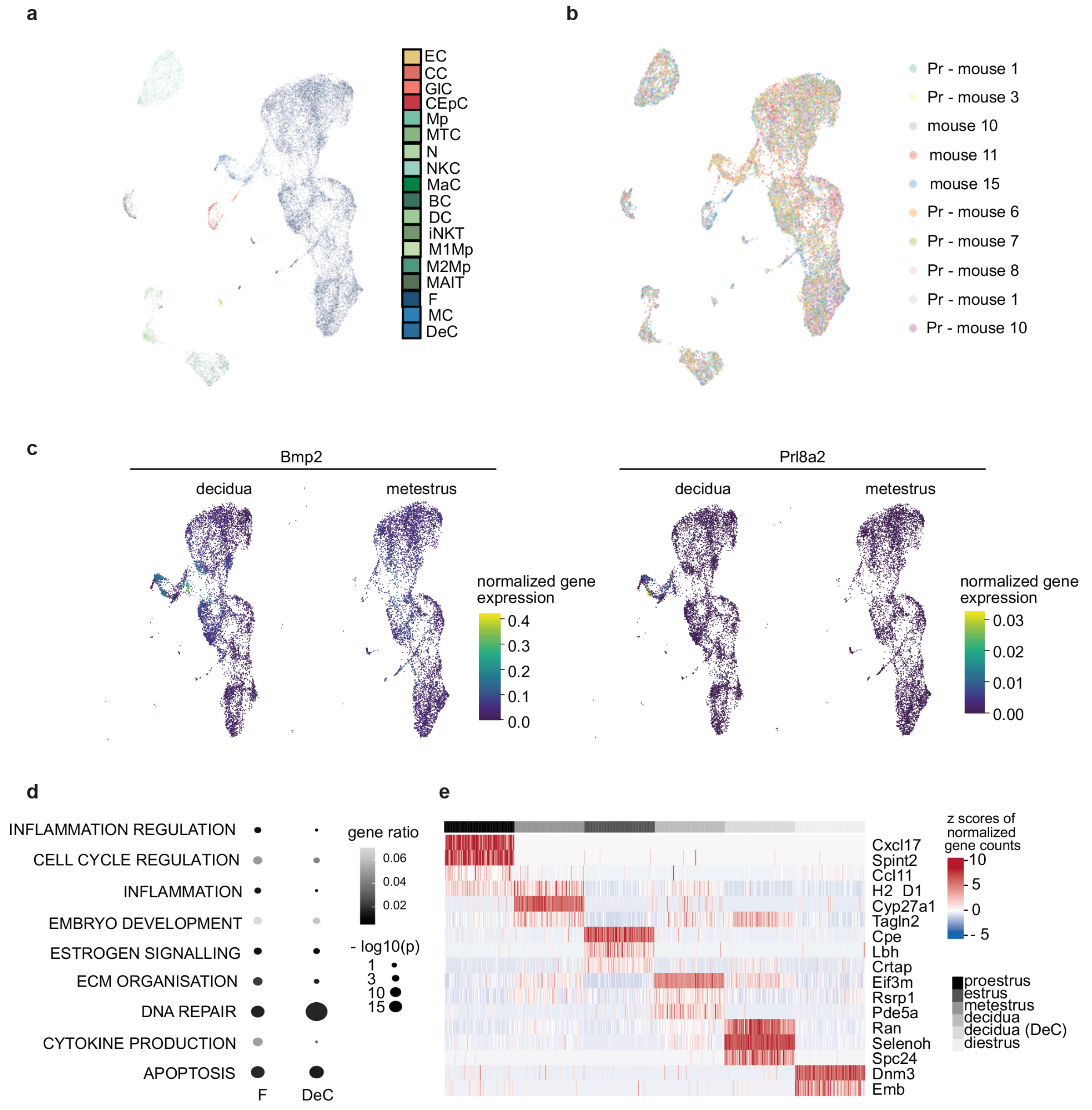


**Figure S8. (a)** UMAP plot of the integration of the pregnant with the metestrus samples. Shown in this panel is a subsample of 19,724 cells. **(b)** UMAP plot of the integration of the pregnant with the metestrus samples colored by individual mice. Uniform mixing of the cells originating from different mice indicates no batch effect present. **(c)** Integrated UMAP plot (see panel a) subsetted to stromal cells and split by condition showing the expression of marker genes of decidualization (*Bmp2* and *Prl8a2*). **(d)** Over-representation analysis of differentially expressed genes between metestrus and pregnant mice in fibroblasts and decidual cells using multilevel generalized linear model (Methods). **(e)** Normalized gene expression of representative differentially expressed genes across the estrus cycle and during decidualization in pregnancy in fibroblasts and decidual cells. Differentially expressed genes were identified using the Mutual Information (MI) approach (Methods).

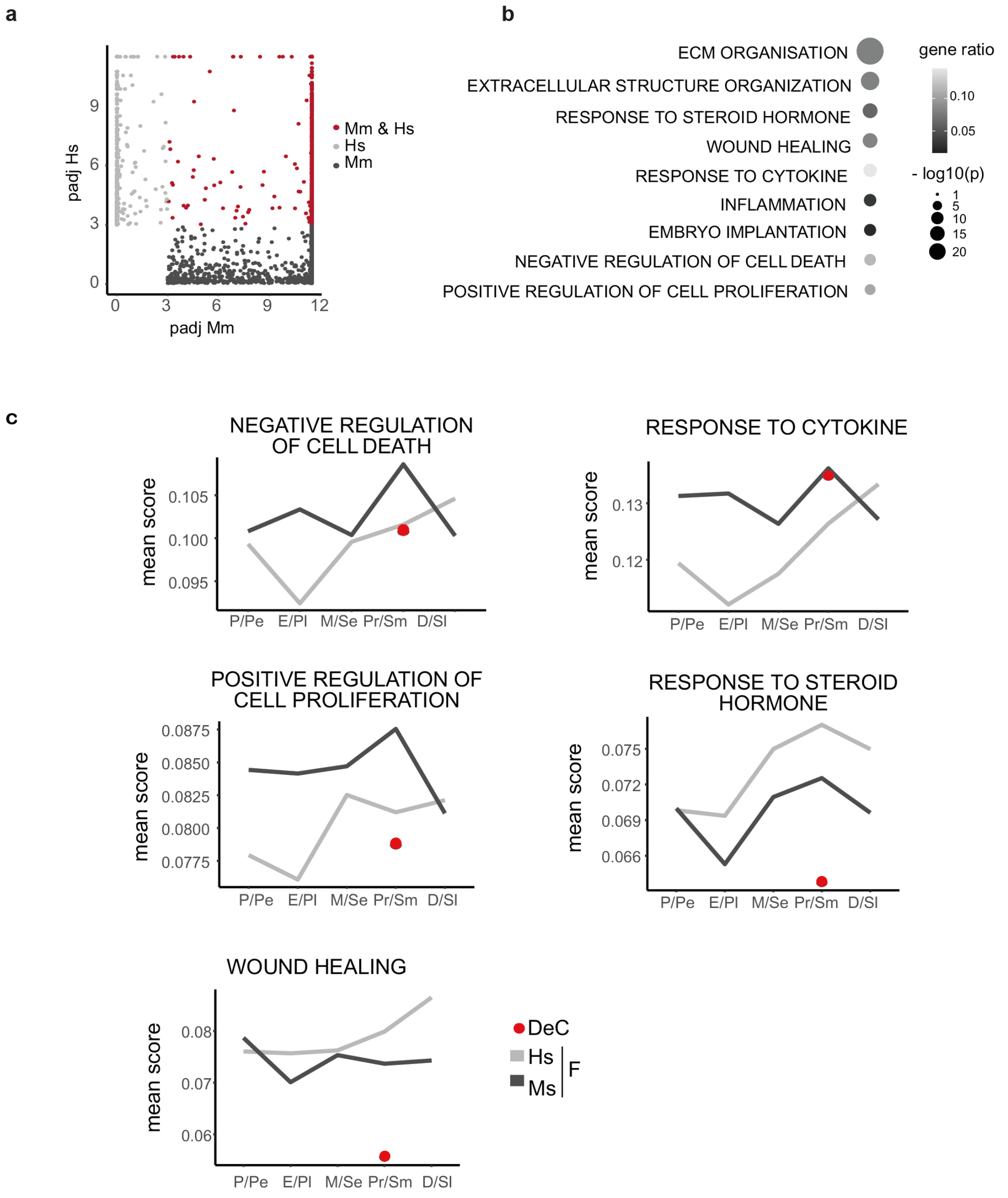


**Figure S9. (a)** Scatter plot of adjusted p-values of mouse and human homologous differentially expressed genes across the cycles calculated using the MI approach. **(b)** ORA analysis of differentially regulated genes that intersected in mouse and human cycles identified using the MI approach (Methods). **(c)** Activity scores of genes in overrepresented pathways (panel b) determined by AUCell and averaged across all mouse and human fibroblasts and mouse decidual cells (Sup. Table S9) in paired cycle phases.

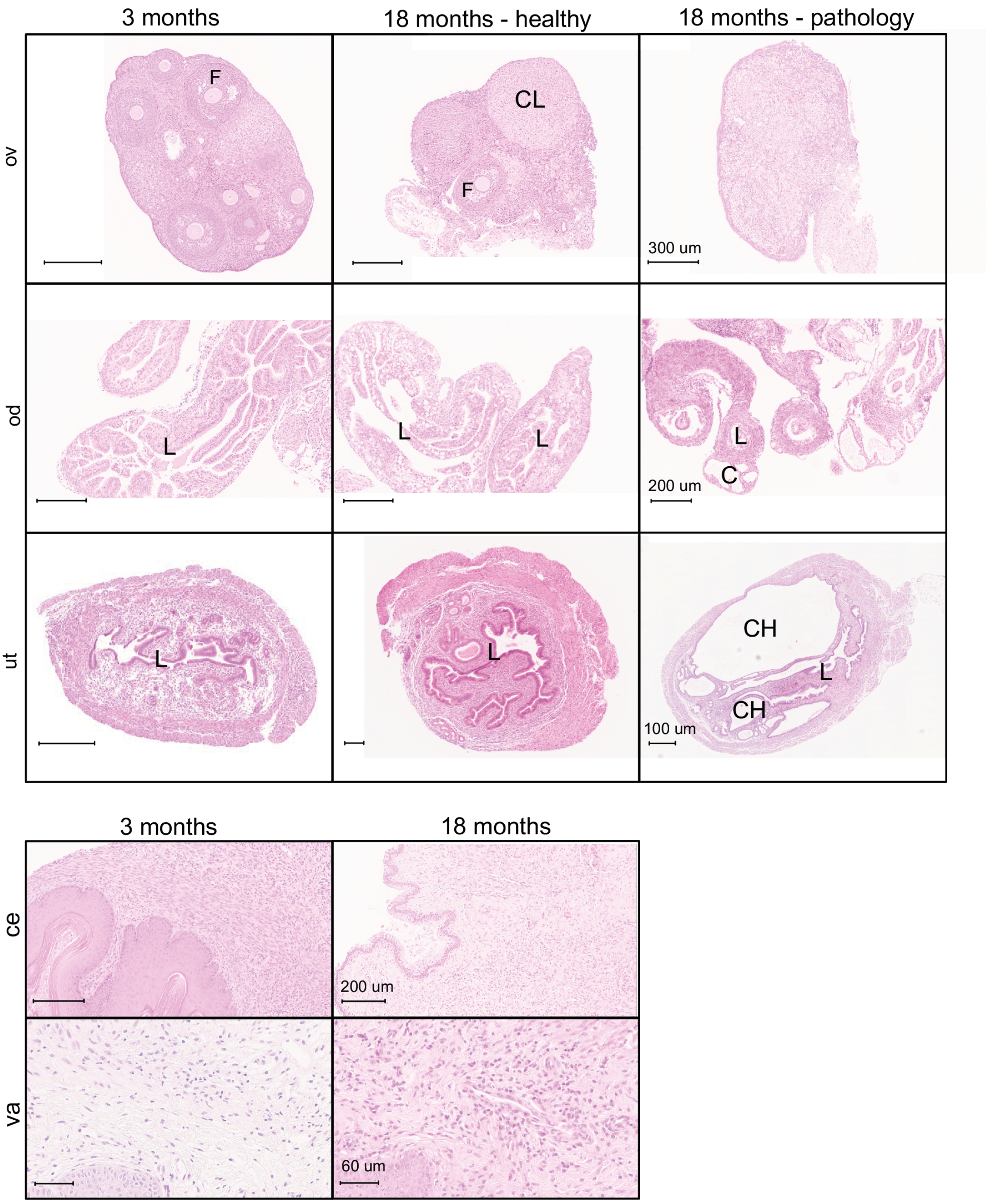


**Figure S10.** Representative H&E staining of young and aged (healthy and pathological) FRT tissues. Pathological ovary is atrophied as evidenced by the absence of follicles and corpus lutea. Follicles and corpus luteum are indicated in young and healthy old with letters F and CL, respectively. In the pathological oviductal sample adenomyosis is visible. L indicates lumen and C cystic dilated glands within the muscle layer. Pathological uterus shows presence of cystic dilated and hyperplastic endometrial glands (CH). Cervical and vaginal tissue show immune cell infiltration in old age samples.

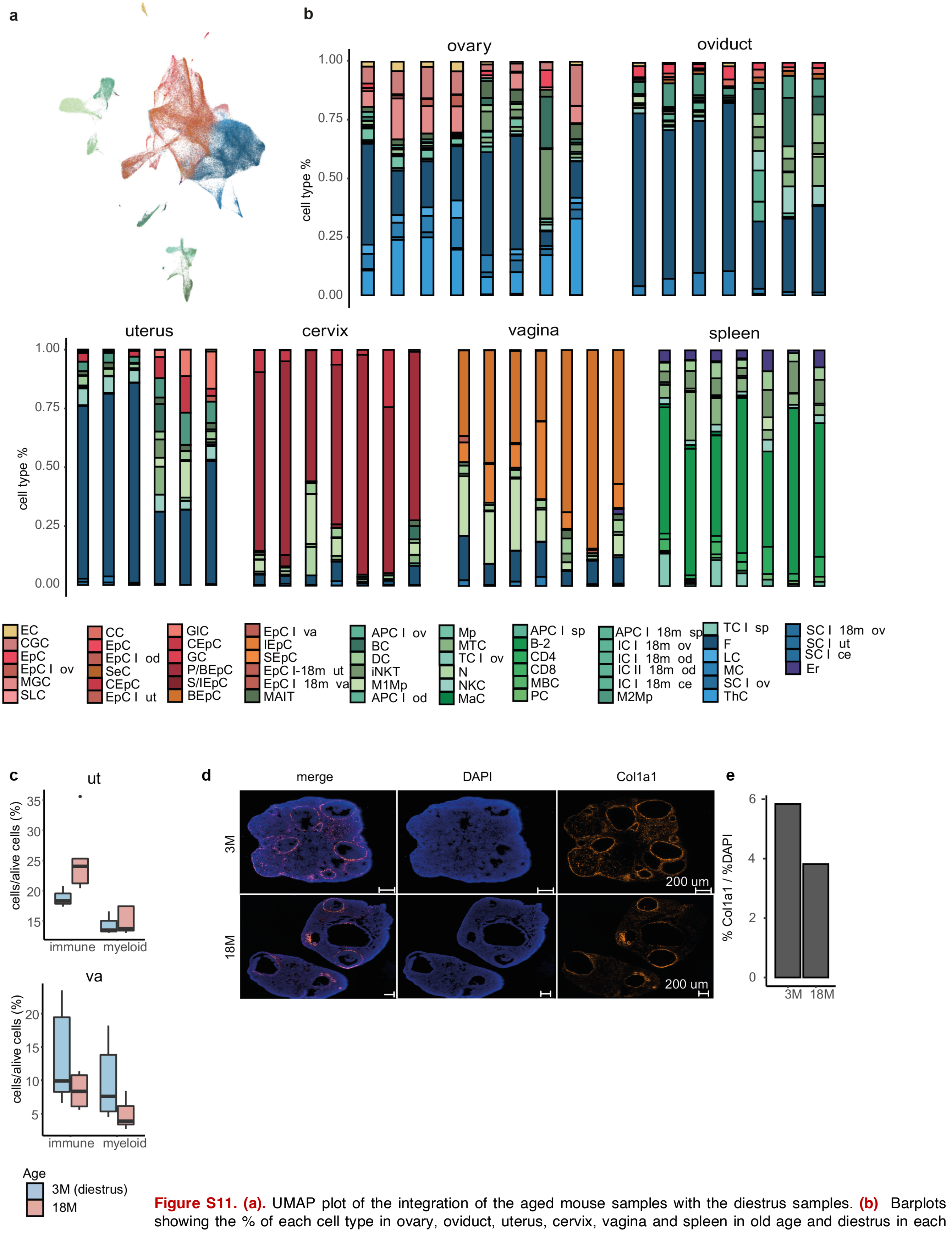


**Figure S11. (a).** UMAP plot of the integration of the aged mouse samples with the diestrus samples. **(b)** Barplots showing the % of each cell type in ovary, oviduct, uterus, cervix, vagina and spleen in old age and diestrus in each biological replicate. **(c)** Quantification of immune and myeloid cell proportions in the uterus and vagina of young and old mice using FACS. **(d)** Expression of Col1a1 in the uterus of young and old mice detected using RNA hybridisation. **(e)** Quantification of Col1a1 signal shown in (d). Col1a1 signal is defined as percent area of total DAPI area. Bar height indicates the average value of two biological replicates.

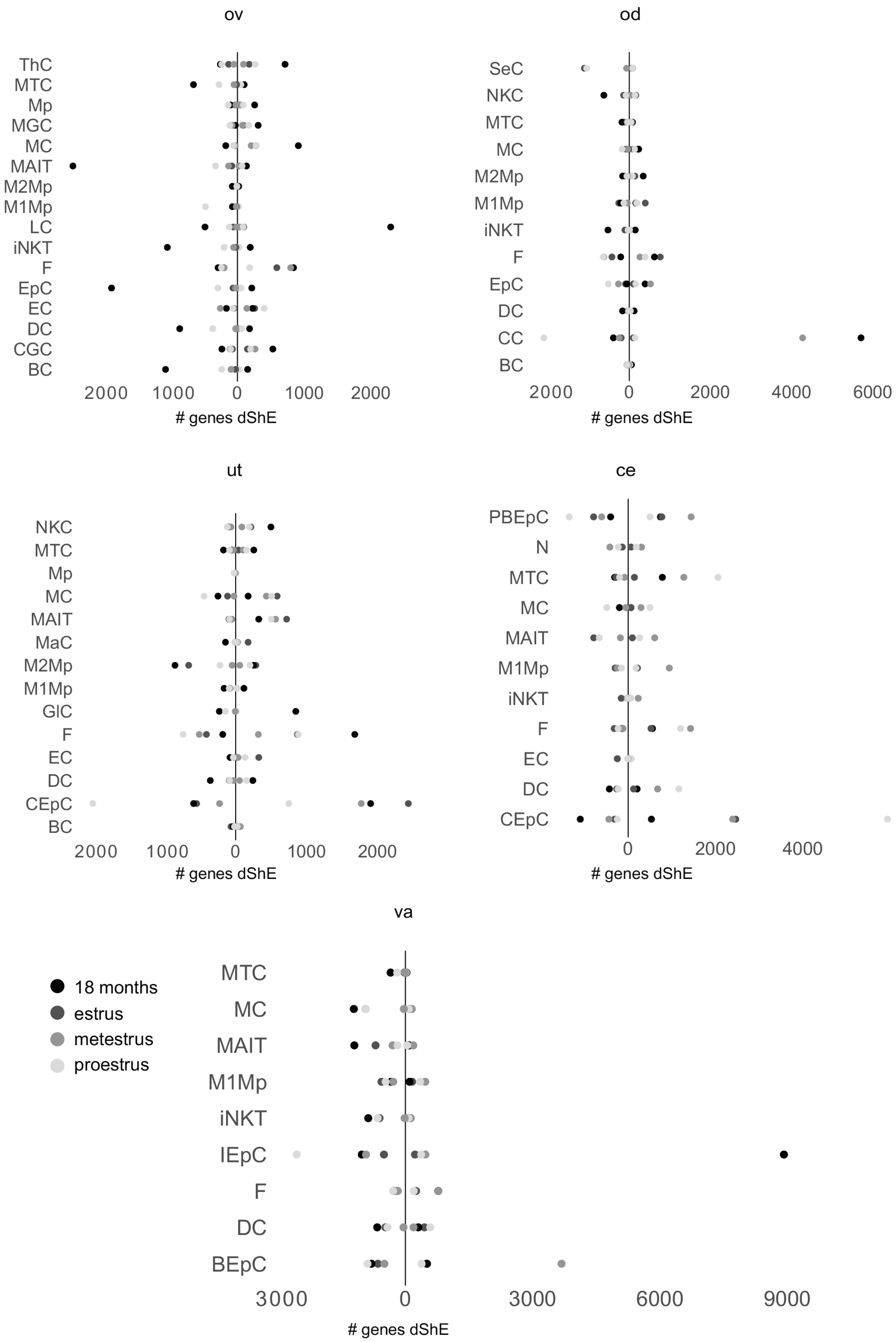


**Figure S12.** Number of genes with differential Shannon entropy (ShE) of all cell types in ovary, oviduct, uterus, cervix and vagina in diestrus compared to other phases of the cycle and old age. Dots on the left side of the y-axis indicate increased entropy relative to diestrus (e.g. higher ShE in old age compared to diestrus) and dots on the right side indicate decreased entropy.

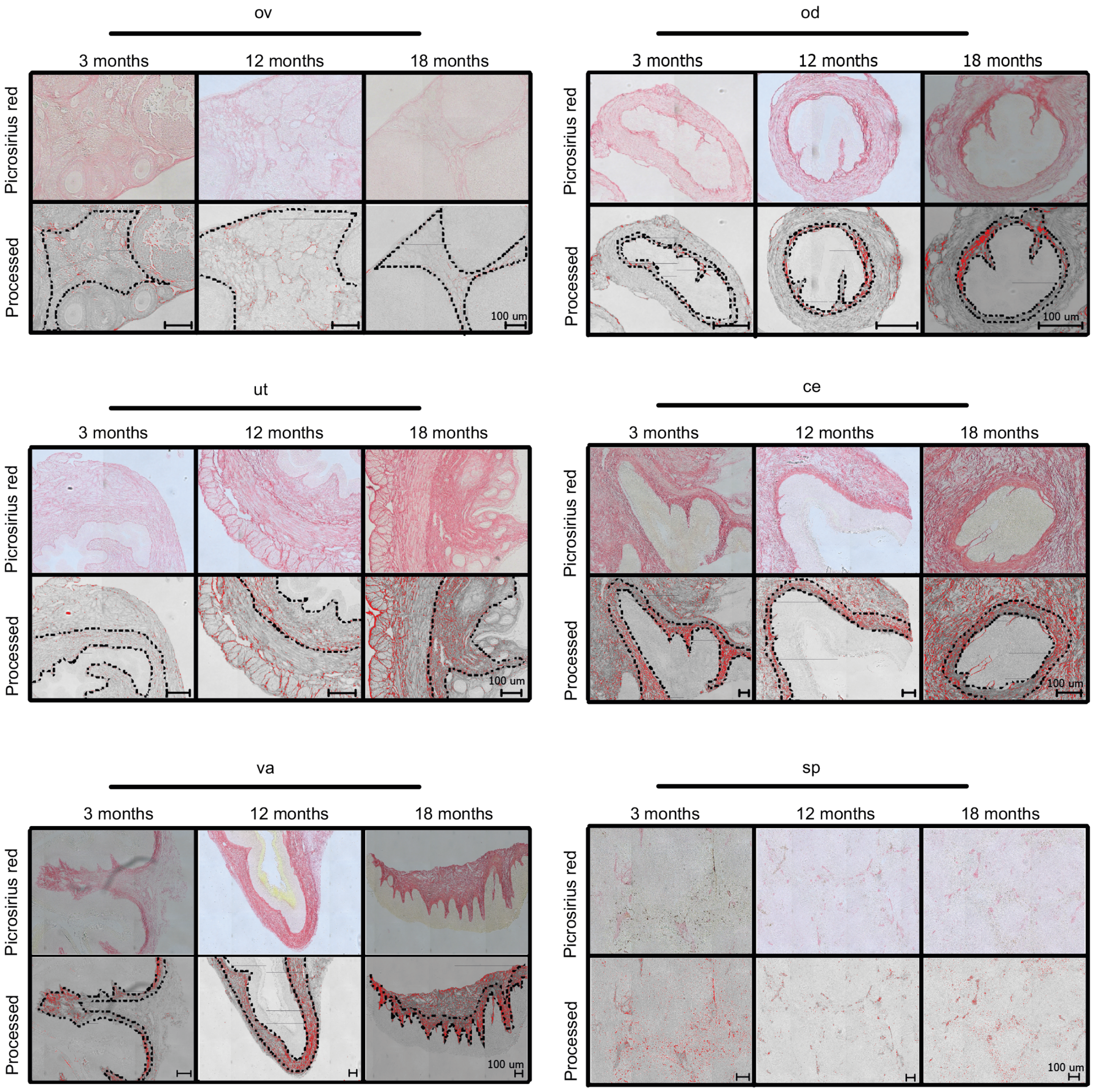


**Figure S13.** % area of stained collagen deposition in ovarian, oviductal, uterine, cervical and vaginal tissue of 3, 12, and 18 month-old mice. For each tissue and time point raw are processed images are shown. In processed images the quantified stromal area is indicated.
